# Disrupted Mitochondrial Copper Homeostasis Promotes Ferroptotic Stress, Senescence and MASLD Progression

**DOI:** 10.64898/2026.06.15.731916

**Authors:** Niansheng Ren, Liuyang Wang, Rajesh K. Dutta, David S. Umbaugh, Qiaojuan Zhang, Seh Hoon Oh, Dennis C. Ko, Ming Song, Anna Mae Diehl, Kuo Du

## Abstract

**Background & Aims:** Systemic metabolic dysfunction promotes degenerative diseases in many organs, including liver and kidney. The liver is a master regulator of systemic metal ion homeostasis. Hepatic copper deficiency is increasingly observed in metabolic dysfunction associated steatotic liver disease (MASLD) and is associated with greater disease severity and poor outcomes. However, mechanisms linking copper dysregulation to MASLD and its co-morbidities remain poorly defined. We investigated whether impaired mitochondrial copper homeostasis contributes to MASLD-related pathobiology and represents a modifiable therapeutic axis.

**Methods & Results:** Using dietary mouse models of MASLD and *in vitro* systems, we found that dietary copper deficiency induces lipotoxicity and suppresses mitochondrial metabolic programs. MASLD livers exhibited marked depletion of copper, impaired cytochrome c oxidase integrity, and bioenergetic failure. Targeted restoration of mitochondrial copper with the copper ionophore elesclomol normalized copper-handling programs, improved mitochondrial function, and suppressed ferroptotic stress, hepatocyte senescence, and fibroinflammatory remodeling. Mechanistically, reduced expression of the mitochondrial copper transporter *SLC25A3* and *MT-CO1* disrupted the SLC25A3–SCO1–MT-CO1–CTR1 axis, limited copper uptake and destabilized copper-iron balance, promoting maladaptive cell fate changes. Across multiple human cohorts and mouse models, copper-iron imbalance tracks with MASLD progression, clinical outcomes, and multiple extrahepatic comorbidities; restoring copper homeostasis in mice with MASLD attenuates both liver and kidney inflammation and fibrosis.

**Conclusions:** Mitochondrial copper deficiency is a mechanistically actionable driver of MASLD that promotes bioenergetic failure, ferroptosis, senescence and fibroinflammatory damage in the liver and other organs. Targeting copper-centered mitochondrial regulation represents a novel biomarker and therapeutic strategy for MASLD and its systemic complications.

## INTRODUCTION

The liver is a master regulator of systemic metabolic homeostasis^1, 2^. Metabolic dysfunction-associated steatotic liver disease (MASLD) is one of the most prevalent chronic liver diseases worldwide and a rapidly increasing cause of cirrhosis and hepatocellular carcinoma (HCC)^3^. Disease progression from steatosis to steatohepatitis and fibrosis reflects the inability of hepatocytes to adapt to sustained metabolic stress, leading to mitochondrial dysfunction, oxidative injury, inflammation, and fibrotic remodeling^4^. Importantly, MASLD rarely occurs in isolation; its clinical course and outcomes are strongly shaped by systemic metabolic comorbidities, including obesity, diabetes, chronic kidney disease and cardiovascular disease^3^. Despite substantial advances in understanding lipid metabolism and fibroinflammatory response, effective disease-modifying therapies remain limited, underscoring the need to identify upstream regulators that integrate metabolic stress with hepatocyte fate decisions and disease progression.

Malnutrition and micronutrient imbalance are highly prevalent in chronic liver disease and are strong, independent predictors of morbidity and mortality^5, 6^. Yet, mechanistic studies in MASLD have largely emphasized macronutrients such as fat, calories, and protein, while deficiencies in trace elements remain comparatively understudied. This represents a critical gap, as micronutrients, including trace metals, exert disproportionate effects on mitochondrial function, redox balance, and cellular stress responses, processes that are fundamental not only to liver injury but also to degeneration of extrahepatic metabolic organs^5, 6^. Thus, micronutrient dysregulation may represent a shared, systemic vulnerability rather than a liver-restricted abnormality. Among trace elements, copper dysregulation has emerged as a distinctive and clinically important feature of MASLD. Copper is an essential trace element required for lipid and glucose metabolism, mitochondrial respiration, antioxidant defense, and iron homeostasis^7–12^. While extreme copper imbalance causes liver injury in rare genetic disorders, converging epidemiologic, human, and experimental data indicate that *acquired* hepatic copper deficiency is common in MASLD and strongly associates with steatohepatitis, fibrosis, metabolic dysfunction, and adverse clinical outcomes^13^. Epidemiologically, Western diets often provide marginal copper, leaving a substantial fraction of adults below recommended intakes^14^. In experimental models, dietary copper restriction induces hepatic steatosis and insulin resistance^13, 15^, while feeding high-sucrose or high-fructose diets further exacerbates copper deficiency, promotes hepatic iron accumulation, and aggravates liver injury^16–18^. Consistent with these observations, a multi-etiology case series of adults with chronic liver disease documented copper deficiency across diverse etiologies, and copper supplementation improved serum copper, ceruloplasmin, and liver function parameters in patients^19^. Together, these findings indicate that copper deficiency in MASLD is prognostically meaningful, at least partially reversible, and may represent a modifiable contributor to disease rather than a passive consequence of liver injury. However, the mechanisms by which copper deficiency promotes MASLD progression remain poorly defined.

We propose that mitochondria act as a central nexus linking copper biology to MASLD pathogenesis because: *i)* mitochondrial dysfunction is a defining feature of MASLD and a key upstream driver of maladaptive hepatocyte responses, including senescence^20, 21^; and *ii)* proper functioning of the mitochondrial electron transport chain (ETC) requires copper to balance the actions of iron^22^. Iron is an obligatory component of the redox active clusters in Complexes I, II and III that ultimately transfer elections to complex IV (cytochrome C oxidase; COX), the copper-dependent terminal enzyme in the ETC that determines whether electrons actually reduce molecular oxygen to generate water^22^. Successful electron transfer drives proton pumping that is required for ATP synthesis by Complex V. Without copper, Complex IV becomes nonfunctional, electrons back up in the ETC, proton pumping fails, ATP synthesis collapses and oxygen consumption drops^22^. Copper also critically regulates iron bioavailability by activating copper-dependent multicopper oxidases such as ceruloplasmin, whose ferroxidase activity is required for iron mobilization and release from hepatocytes^23, 24^. Copper deficiency, therefore, promotes excessive retention of iron (Fe) in an electron-enriched environment that favors both production of reactive oxygen species and accumulation of Fe^2+^ ions, potent catalysts of membrane lipid peroxidation that promotes ferroptosis, an iron-dependent cell death program that has been implicated in MASLD pathogenesis^25^. In addition, copper deficiency may amplify susceptibility to ferroptosis and other cell death programs by limiting the activity of mitochondrial superoxide dismutase, another copper-dependent enzyme involved in mitochondrial redox control^22^. Because the liver is the principal source of circulating copper- and iron-regulatory proteins, as well as a master regulator of systemic metabolism^1, 2^, dysregulated hepatic copper-iron homeostasis has the potential to propagate metabolic stress and organ dysfunction beyond the liver. Despite this, how mitochondrial copper-iron imbalance shapes hepatocyte fate decisions, systemic vulnerability, and MASLD progression has remained underexplored.

Here, we address this critical gap by systematically investigating the role of dysregulated copper homeostasis in MASLD. Using complementary dietary mouse models of steatohepatitis, hepatocyte-based *in vitro* systems, and integrative transcriptomic analyses, we examine how alterations in copper handling, and its coupling to iron metabolism, influence mitochondrial bioenergetic function, hepatocyte stress responses, and disease progression. We further leverage bulk and single-cell transcriptomic datasets from human livers and multiple organs to assess whether signatures of mitochondrial copper sufficiency and iron imbalance associate with MASLD severity, therapeutic response, hepatocellular carcinoma risk, and extrahepatic metabolic comorbidities that drive poor clinical outcomes. Together, these studies aim to define mitochondrial copper-iron regulation as a previously underappreciated, shared metabolic axis that links disrupted trace metal homeostasis to cellular vulnerability, organ degeneration, and MASLD pathogenesis.

## RESULTS

### Dietary copper deficiency induces MASLD-like features and suppresses mitochondrial programs

To define the molecular consequences of copper deficiency in metabolic liver disease, we reanalyzed liver transcriptomes from rats fed a copper-deficient AIN-76A diet (<0.3 mg/kg Cu) or a copper-adequate diet (125 mg/kg Cu) for 12 weeks^18^ **(Supplementary Fig. 1A)**. Gene set enrichment analysis (GSEA) demonstrated broad pathway remodeling in copper-deficient livers, characterized by depletion of core hepatic metabolic programs (e.g., *bile acid metabolism*; *xenobiotic metabolism)* but enrichment of inflammatory and fibrotic pathways (e.g., *inflammatory response*; *TNF*α *signaling via NF-*κ*B*; *ECM receptor interaction*) **(Supplementary Figs. 1B, C)**. Notably, mitochondrial metabolic programs were markedly suppressed (e.g., *fatty acid metabolism; oxidative phosphorylation; and lipid oxidation*) **(Supplementary Figs. 1B, C)**. Consistent with these transcriptomic changes, prior ultrastructural studies in copper-deficient liver also demonstrate progressive mitochondrial enlargement and megamitochondria-like morphology^26–28^, features linked to impaired respiratory capacity and heightened oxidative stress. Collectively, these data indicate that dietary copper deficiency is sufficient to induce mitochondrial dysfunction and transcriptional remodeling that mediate MASLD pathogenesis.

### Elesclomol increases mitochondrial copper, normalizes copper-handling programs, and protects against MASLD in preclinical models

Copper is indispensable for mitochondrial respiration through cytochrome c oxidase (Complex IV)^29–31^. During copper scarcity, cells prioritize mitochondrial delivery to preserve bioenergetic capacity^32^. To test whether mitochondrial copper can be selectively augmented *in vivo*, chow-fed mice were treated with the mitochondrial copper ionophore elesclomol^33–36^ (10 mg/kg, oral gavage), followed by inductively coupled plasma mass spectrometry (ICP-MS)-based metallomic analysis **(Fig. 1A)**. Elesclomol robustly increased mitochondrial copper without altering total hepatic copper levels **(Fig. 1A)**, establishing the feasibility of organelle-targeted copper repletion and validating elesclomol as a tool to interrogate the functional consequences of mitochondrial copper sufficiency.

**Fig. 1.**
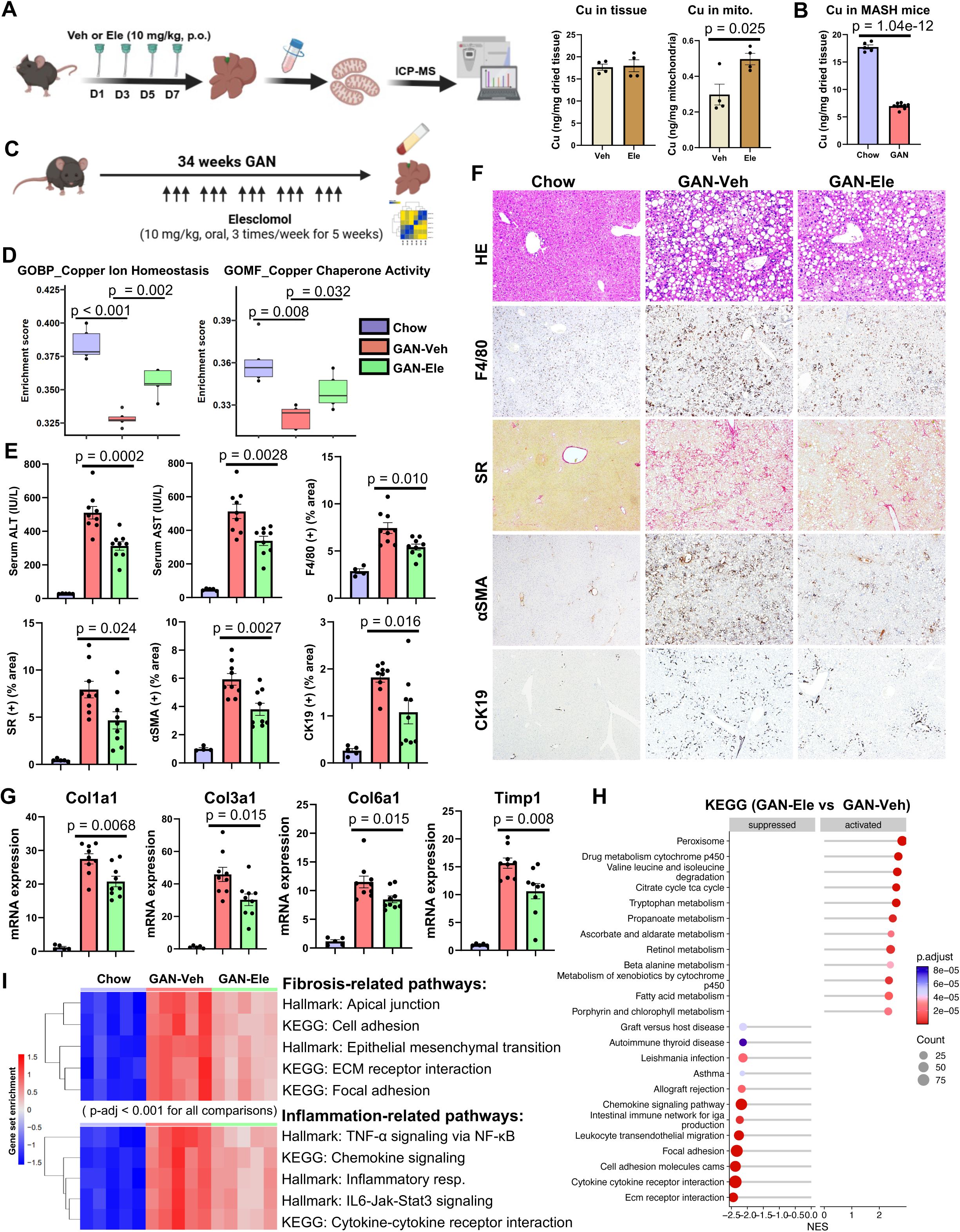
Mitochondrial copper ionophore elesclomol ameliorates MASLD in GAN diet-fed mice. **(A)** Experimental design for short-term elesclomol treatment and copper quantification. Mice received vehicle or elesclomol (10 mg/kg, oral gavage) for 7 days, followed by liver harvest, mitochondrial isolation, and copper measurement by ICP-MS (mean ± SEM, n = 4 per group; Student’s *t* test). **(B)** Hepatic copper levels in mice fed with chow diet or Gubra Amylin NASH (GAN) diet for 34 weeks. **(C)** Experimental design for long-term MASLD studies. Mice were fed a GAN diet for 34 weeks and treated with vehicle or elesclomol during the final 5 weeks (10 mg/kg, oral gavage, three times per week; chow n = 4, GAN-Veh and GAN-Ele n = 9). **(D)** Gene set enrichment analysis (GSEA) showing enrichment scores for copper homeostasis and copper chaperone activity pathways. **(E)** Serum alanine aminotransferase (ALT), aspartate aminotransferase (AST), and quantitative analysis of positively stained areas shown in F. **(F)** Representative liver histology showing hematoxylin and eosin (H&E), Picrosirius Red (SR), F4/80, α-SMA, and CK19 staining. **(G)** Relative hepatic mRNA expression of fibrosis-related genes assessed by qRT-PCR. **(H)** KEGG pathway enrichment analysis comparing GAN-Ele versus GAN-Veh livers. **(I)** Heatmap showing Hallmark and KEGG fibroinflammatory pathway enrichment across experimental groups. Data are shown as mean ± SEM. Statistical significance in **A-G** was determined by Student t-test (2 groups) or one-way ANOVA with post hoc testing (3 groups) as indicated. Enrichment analyses in **H-I** were performed using permutation testing with Benjamini–Hochberg correction for multiple comparisons.

We next examined whether hepatic copper is depleted in MASH (Metabolic dysfunction-Associated Steatohepatitis; severe MASLD with inflammation and cellular damage) and whether this deficit is reversible. In mice fed the GAN (Gubra-Amylin NASH diet) diet for 34 weeks to induce MASH, total hepatic copper was reduced by more than 60% relative to chow-fed controls **(Fig. 1B)**, confirming a profound copper deficit in advanced MASLD. To evaluate therapeutic efficacy, mice were maintained on the GAN diet for 34 weeks and randomized to vehicle or elesclomol treatment for the final 5 weeks **(Fig. 1C)**. Transcriptomic profiling revealed that the GAN diet markedly disrupted copper homeostasis pathways and copper-chaperone activity, as assessed by GSEA of copper-handling programs **(Fig. 1D)**. Strikingly, these pathways were significantly restored by elesclomol treatment **(Fig. 1D)**, indicating effective correction of copper-dependent molecular networks in diseased liver. Restoration of copper homeostasis was accompanied by significant improvements in biochemical indices of liver injury, including reductions in serum ALT and AST **(Fig. 1E)**. Importantly, these beneficial effects were not attributable to changes in body weight, which was comparable between GAN-vehicle and GAN-elesclomol groups **(Supplementary Fig. 2A)**. Instead, elesclomol selectively reduced diet-induced hepatomegaly and improved metabolic and cholestatic parameters, including reductions in serum cholesterol and bilirubin **(Supplementary Figs. 2A, B)**. Histopathologic analyses further demonstrated that restoration of mitochondrial copper markedly attenuated MASLD features, as evidenced by reduced steatosis on H&E staining, decreased macrophage infiltration (F4/80), diminished collagen deposition (Sirius Red), reduced hepatic stellate cell activation (αSMA), and attenuated ductular reaction (CK19) **(Figs. 1E, F)**. These structural improvements were accompanied by significant downregulation of key extracellular matrix and fibrogenic genes, including *Col1a1*, *Col3a1*, *Col6a1*, and *Timp1* **(Fig. 1G).** Pathway analyses further revealed coordinated improvement of core hepatic metabolic programs (e.g., *bile acid metabolism; fatty acid metabolism; cytochrome P450-mediated drug metabolism*) together with suppression of fibrosis- and inflammation-related signaling pathways (e.g., *extracellular matrix organization; TNF*α*/NF-*κ*B signaling; cytokine–cytokine receptor interaction*) with elesclomol treatment **(Figs. 1H, I; Supplementary Figs. 2C, D)**.

To assess generalizability across MASLD models, we repeated the intervention in choline-deficient, amino acid–defined high-fat diet (CDAHFD)-fed mice **(Fig. 2A)**. Consistent with the GAN model, elesclomol treatment reduced hepatocellular injury and attenuated histologic features of MASLD, including steatohepatitis and fibrosis **(Figs. 2B-E)**. Transcriptomic analyses similarly demonstrated coordinated suppression of inflammation- and fibrosis-related pathways and partial restoration of metabolic programs **(Figs. 2F-H; Supplementary Figs. 3A, B)**. Together with results in the GAN diet model, these data demonstrate that targeted restoration of mitochondrial copper consistently ameliorates hepatocellular injury, suppresses fibroinflammatory remodeling, and improves disease-associated transcriptomic signatures across MASLD models, providing strong cross-model therapeutic proof of concept.

**Fig. 2.**
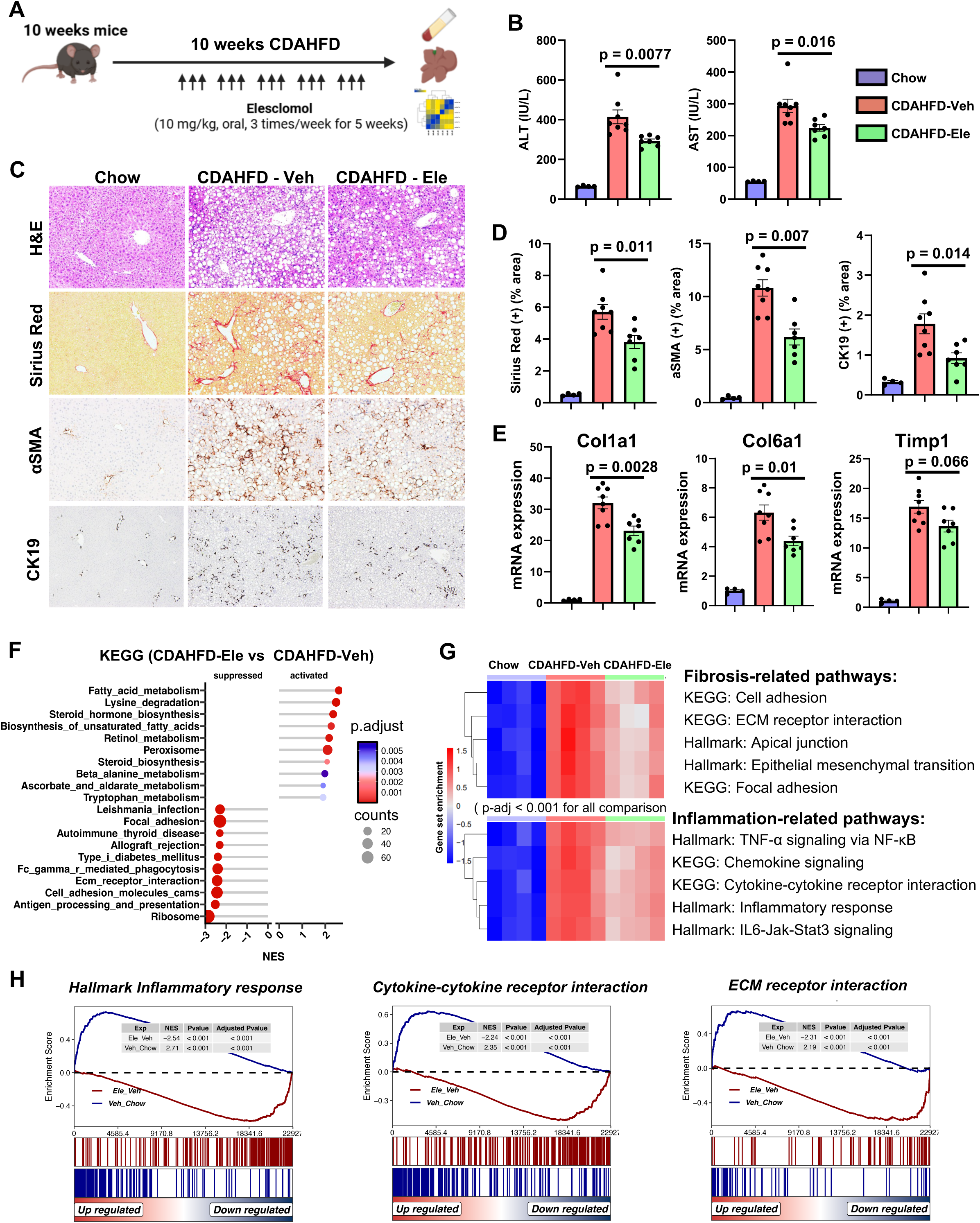
Elesclomol ameliorates MASLD in CDAHFD-fed mice. **(A)** Experimental design. Ten-week-old mice were fed a choline-deficient, L-amino acid–defined high-fat diet (CDAHFD) for 10 weeks and treated with vehicle or elesclomol (10 mg/kg, oral gavage, three times per week) during the final 5 weeks. **(B)** Serum alanine aminotransferase (ALT) and aspartate aminotransferase (AST) levels. **(C)** Representative liver histology showing hematoxylin and eosin (H&E), Picrosirius Red, α-smooth muscle actin (α-SMA), CK19 staining and **(D)** corresponding quantification of positively-stained areas. **(E)** Relative hepatic mRNA expression of fibrosis-related genes assessed by qRT-PCR. **(F)** KEGG pathway enrichment analysis comparing CDAHFD-Ele versus CDAHFD-Veh livers. **(G)** Heatmap showing enrichment of fibrosis- and inflammation-related Hallmark and KEGG pathways across experimental groups. **(H)** Representative GSEA enrichment plots for selected Hallmark and KEGG pathways. Data are shown as mean ± SEM; statistical significance in B-E was determined by one-way ANOVA with post hoc testing as indicated. *P* values in GSEA were calculated using permutation testing and adjusted for multiple comparisons using the Benjamini-Hochberg method.

### Restoring copper homeostasis improves mitochondrial bioenergetics

Copper supports mitochondrial respiration and thereby, metabolic and energy homeostasis, primarily via its role as an essential co-factor for cytochrome c oxidase activity (Complex IV) ^29–31^. Because hepatic copper is depleted in MASLD, we asked whether mitochondrial bioenergetic programs are functionally impaired and whether targeted copper restoration can normalize them.

In both GAN- and CDAHFD-fed mice, transcriptome analyses revealed strong suppression of oxidative phosphorylation, the tricarboxylic acid (TCA) cycle, and fatty acid metabolism **(Figs. 3A, B)**. These transcriptional defects were accompanied by reduced abundance of key electron transport chain (ETC) proteins spanning multiple complexes, including MT-CO1 (Complex IV), ATP5B (Complex V), UQCRC2 (Complex III), and NDUFB8 (Complex I) **(Figs. 3C, D**). We focused on MT-CO1 because it is a copper-dependent Complex IV subunit and a direct marker of cytochrome c oxidase competence^29–31^. Elesclomol has previously been shown to rescue lethality and neuropathology in Menkes disease mice by delivering copper to mitochondria and restoring cytochrome c oxidase in brain tissue^33–36^. Consistent with this mechanism, elesclomol treatment increased MT-CO1 in MASLD livers, together with coordinated restoration of other ETC subunits, indicating recovery of Complex IV-driven respiratory capacity **(Figs. 3C, D**). In contrast, SOD1, a copper enzyme that is predominantly cytosolic with a small pool in the mitochondrial intermembrane space^37^, remained suppressed and was not restored by elesclomol **(Supplementary Fig. 4A, B)**, supporting a mitochondria-targeted, Complex IV-centric mechanism of action. We also assessed ceruloplasmin (CP), the major hepatocyte-derived copper carrier with ferroxidase activity^38^. Although CP has been reported to be downregulated in MASLD as a potential compensatory response^39^, CP protein abundance was increased in GAN-fed mice and was not significantly altered by elesclomol treatment **(Supplementary Figs. 4A, B)**, indicating that therapeutic benefit occurs independently of CP modulation.

**Fig. 3.**
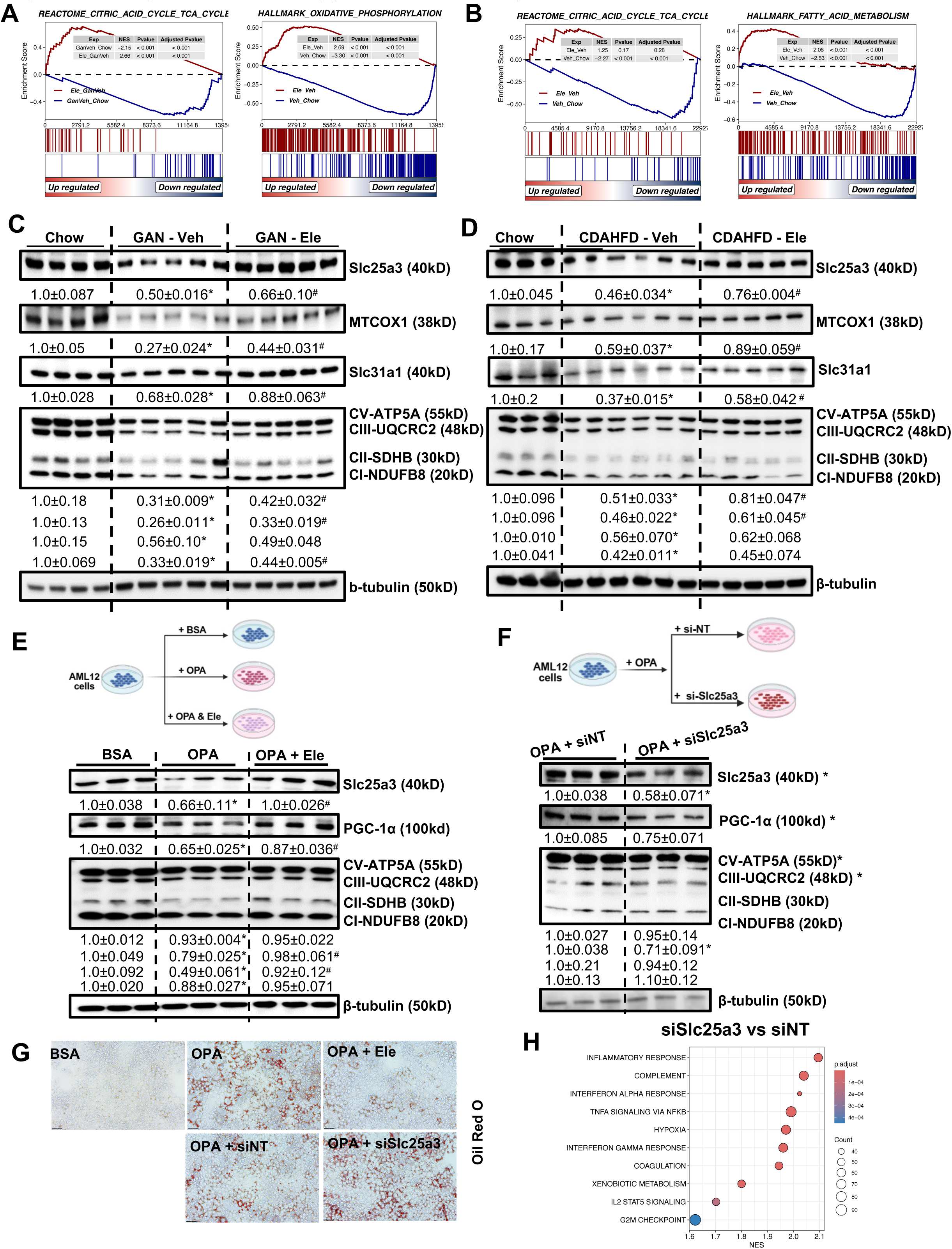
Restoring mitochondrial copper homeostasis improves mitochondrial function in MASH. **(A, B)** GSEA of mitochondrial bioenergetic pathways in livers from (A) GAN- and (B) CDAHFD-fed mice. **(C, D)** Immunoblot analysis of copper transporters and mitochondrial respiratory chain proteins in livers from chow-, GAN-, and CDAHFD-fed mice treated with vehicle or elesclomol, with β-tubulin as a loading control. **(E)** Immunoblot analysis of copper transporters and mitochondrial respiratory chain proteins in AML12 hepatocytes treated with BSA control, oleate/palmitate (OPA), or OPA plus elesclomol. **(F)** Immunoblot analysis of AML12 hepatocytes treated with OPA following transfection with non-targeting siRNA (siNT) or Slc25a3 siRNA. **(G)** Representative Oil Red O staining of AML12 hepatocytes under the indicated conditions. **(H)** Pathway analysis of differentially expressed genes in Slc25a3-deficient AML12 hepatocytes compared with siNT controls. Densitometric quantification is shown below each blot. Data are shown as mean ± SEM. \**p* < 0.05 vs. chow; ^#^*p* < 0.05 vs. vehicle. GSEA *P* values were calculated by permutation testing and adjusted for multiple comparisons using the Benjamini-Hochberg method.

At the level of copper transport, SLC25A3, the principal mitochondrial copper carrier required for inner-membrane delivery and COX metalation and assembly^40, 41^, was reduced in MASLD liver but partially restored by elesclomol **(Figs. 3C, D)**. SLC31A1 (CTR1), the high-affinity plasma-membrane copper importer^42^, was also reduced in MASLD livers, providing a parsimonious explanation for lower total hepatic copper; its partial recovery **(Figs. 3C, D)** helps explain why elesclomol normalizes copper-handling programs and homeostasis **(Fig. 1B)**. Prior work shows that the COX assembly factor SCO1 sustains CTR1 stability^43–45^. When COX assembly falters, SCO1-dependent signaling promotes proteasomal degradation of CTR1, reducing cellular copper entry and further impairing COX biogenesis^43–45^. Together, these data establish a mitochondria-to-plasma membrane feedback loop that amplifies copper scarcity when Complex IV is compromised. By delivering copper to mitochondria, elesclomol functionally bypasses CTR1 limitation, restores COX/MT-CO1, stabilizes respiratory programs, and improves hepatocyte bioenergetics.

To test mechanism directly, we employed an *in vitro* MASLD model in which lipotoxicity was induced by oleic plus palmitic acid (OPA). Consistent with *in vivo* findings, OPA exposure in AML12 hepatocytes lowered SLC25A3 abundance, reduced PGC-1α (master regulator of mitochondrial biogenesis) and respiratory-chain proteins, and increased cellular lipid accumulation **(Figs. 3E, G)**. Co-treatment with elesclomol restored PGC-1α expression, increased ETC subunit abundance including MT-CO1, and reduced intracellular lipid accumulation **(Figs. 3E, G)**. Conversely, siRNA-mediated knockdown of *Slc25a3* during OPA exposure further suppressed respiratory proteins and markedly increased lipid accumulation compared with non-targeting controls **(Figs. 3F, G)**. Transcriptomic analyses further revealed that *Slc25a3* deficiency robustly induced epithelial-mesenchymal transition (EMT) programs together with pro-inflammatory and pro-fibrogenic gene signatures in hepatocytes **(Fig. 3H)**. Together, these data demonstrate that SLC25A3-dependent mitochondrial copper import is required for hepatocyte bioenergetic resilience under lipotoxic stress and position SLC25A3 dysfunction within the SCO1-MTCO1-CTR1 axis as a proximal driver of impaired COX assembly, diminished oxidative capacity, and lipotoxic injury in MASLD.

### Restoring mitochondrial copper homeostasis suppresses ferroptotic stress and hepatocyte senescence in MASLD liver

Converging evidence indicates that both ferroptosis and cellular senescence are potentiated by mitochondrial dysfunction and by disturbances in cellular metal handling^46, 47^. Our group has previously shown that aging heightens hepatocyte susceptibility to ferroptotic stress, worsens MASLD trajectory, and that ferroptosis inhibition partially reverts the aging liver transcriptome toward a more youthful state while attenuating injury^25, 48^. We further demonstrated that hepatocyte senescence is a proximal driver of microenvironmental remodeling in chronic liver disease, and that mitochondrial oxidant stress and bioenergetic failure are key upstream triggers of the senescent hepatocyte phenotype^49–52^. Building on these findings, and on the central role of copper in mitochondrial respiration through cytochrome c oxidase, we hypothesized that restoring mitochondrial copper sufficiency would improve bioenergetics and thereby interrupt feed-forward circuits that promote lipid peroxidation, ferroptotic stress, and senescence.

Guided by this premise, we tested whether targeted mitochondrial copper repletion attenuates ferroptosis and senescence *in vivo*. In both GAN- and CDAHFD-fed mice, elesclomol treatment significantly reduced enrichment scores for hepatic iron accumulation and ferroptosis driver signatures **(Figs. 4A, B)**. Notably, ICP-MS analysis revealed that elesclomol reduced hepatic iron content without increasing total hepatic copper levels **(Supplementary Fig. 5A)**. Consistent with reduced ferroptotic stress, lipid peroxidation was markedly decreased, as evidenced by lower malondialdehyde (MDA) levels in liver lysates from elesclomol-treated mice across both dietary models **(Figs. 4C, D)**. Histologic and molecular analyses revealed fewer senescent (p21-positive) hepatocytes and decreased expression of p16, p21 and senescence-associated signatures, including AHGS^25^, SHGS^51^, SenMayo^53^ and SenNet **(Figs. 6E-J; Supplementary Fig. 5B)**.

**Fig. 4.**
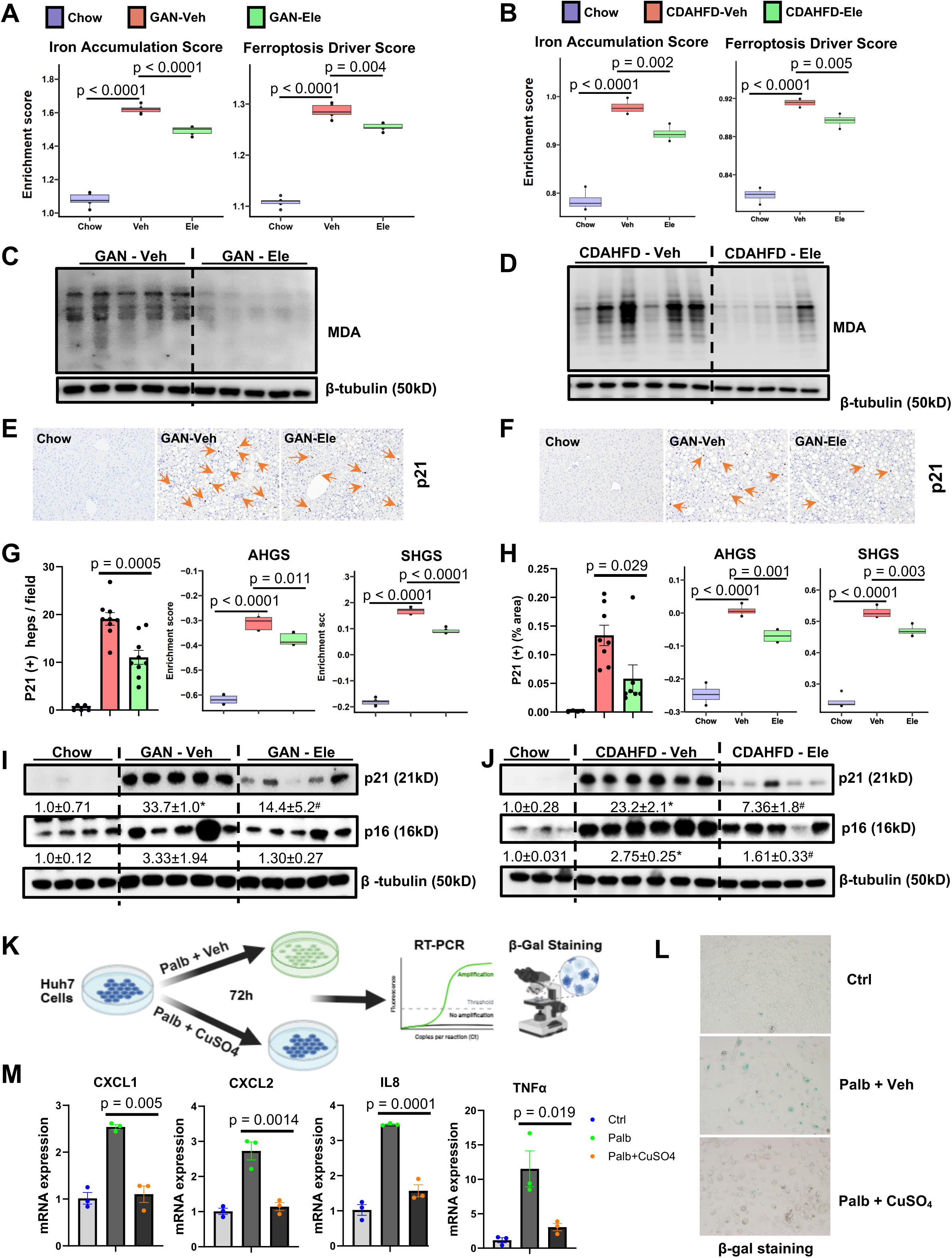
Restoring mitochondrial copper homeostasis suppresses ferroptotic stress and hepatocyte senescence. **(A, B)** Enrichment scores for iron accumulation and ferroptosis driver gene signatures in livers from chow-, GAN-, and CDAHFD-fed mice treated with vehicle or elesclomol. **(C, D)** Immunoblot analysis of malondialdehyde (MDA)-modified proteins in GAN and CDAHFD livers, with β-tubulin as a loading control. **(E, F)** Representative immunohistochemical staining for p21 in GAN and CDAHFD liver sections (arrows indicate positive nuclei). **(G, H)** Quantification of p21-positive area and enrichment scores for aging hepatocyte gene signature (AHGS) and senescent hepatocyte gene signature (SHGS). **(I, J)** Immunoblot analysis of p21 and p16 protein expression in GAN and CDAHFD livers; densitometric quantification is shown below. **(K)** Experimental design for induction of senescence in Huh7 cells using palbociclib (Palb) with or without copper supplementation (CuSO₄). **(L)** Representative senescence-associated β-galactosidase staining in Huh7 cells under indicated conditions. **(M)** Relative mRNA expression of senescence-associated secretory phenotype (SASP) factors assessed by qRT-PCR. Data are shown as mean ± SEM; statistical significance was determined by one-way ANOVA with post hoc testing as indicated. *P* values in GSEA were calculated using permutation testing and adjusted for multiple comparisons using the Benjamini-Hochberg method.

Because high concentrations of the elesclomol-copper complex can induce cuproptosis in cancer contexts^36^, we explicitly evaluated whether this pathway was engaged in MASLD livers.

Notably, elesclomol treatment did not increase expression of cuproptosis marker gene signature in either dietary model **(Supplementary Fig. 5C)**. Moreover, elesclomol consistently reduced liver injury and disease severity **(Figs. 1, 2)**, indicating that therapeutic mitochondrial copper repletion occurs without triggering cuproptotic cell death in this context.

To determine whether copper sufficiency directly modulates senescence outputs in human hepatocytes, we induced senescence in Huh7 cells using the CDK4/6 inhibitor palbociclibb^51^. Copper supplementation with CuSO₄ significantly reduced the proportion of β-galactosidase-positive cells and suppressed expression of canonical Senescence-Associated Secretory Phenotype (SASP) factors, including *CXCL1*, *CXCL2*, *IL8*, and *TNF*α **(Figs. 4K-M)**. These data demonstrate that copper availability directly regulates senescence and SASP activity in hepatocytes, independent of systemic effects. Collectively, these findings demonstrate that restoring mitochondrial copper homeostasis suppresses ferroptotic stress and hepatocyte senescence in both *in vivo* and *in vitro* models. By targeting an upstream micronutrient control point, mitochondrial copper repletion disrupts key ferroptosis-senescence circuits that drive fibroinflammatory remodeling and disease progression in MASLD.

### Copper-iron imbalance tracks with MASLD progression, regression, and clinical outcomes

Building on our experimental data showing that mitochondrial copper deficiency impairs Complex IV-dependent bioenergetics while iron overload promotes lipid peroxidation and ferroptotic stress, we developed a transcriptome-based Cu-Mito Score (CMS) to quantify mitochondrial copper sufficiency. CMS was derived from genes involved in copper uptake, mitochondrial copper transport, and COX biogenesis^54^ (e.g., SLC31A1, SLC25A3, COX subunits, copper chaperones) and was benchmarked against a validated Iron Accumulation Score (IAS) capturing iron overload-associated programs^55^ **(Figs. 5A; Supplementary Fig. 6A)**. To establish the biological relevance of CMS at the cellular level, we examined its enrichment in hepatocytes using single-cell RNA-seq datasets from MASLD patients. Compared with controls, hepatocytes from MASLD livers exhibited a marked reduction in CMS and a concomitant increase in IAS enrichment **(Fig. 5B; Supplementary Fig. 6B)**. This is accompanied by decreased expression of *SLC25A3* and *MT-CO1* and dysregulated copper homeostatic pathways *(e.g., copper chaperone activity; copper iron binding)* **(Fig. 5B; Supplementary Fig. 6C)**, indicating impaired mitochondrial copper transport, compromised COX integrity and dysregulated cellular copper homeostasis at single-cell resolution. Consistent with these transcriptomic changes, immunohistochemical analyses demonstrated a progressive loss of MT-CO1 protein with advancing fibrosis stage in human MASLD liver biopsies **(Supplementary Fig. 6E)**.

**Fig. 5.**
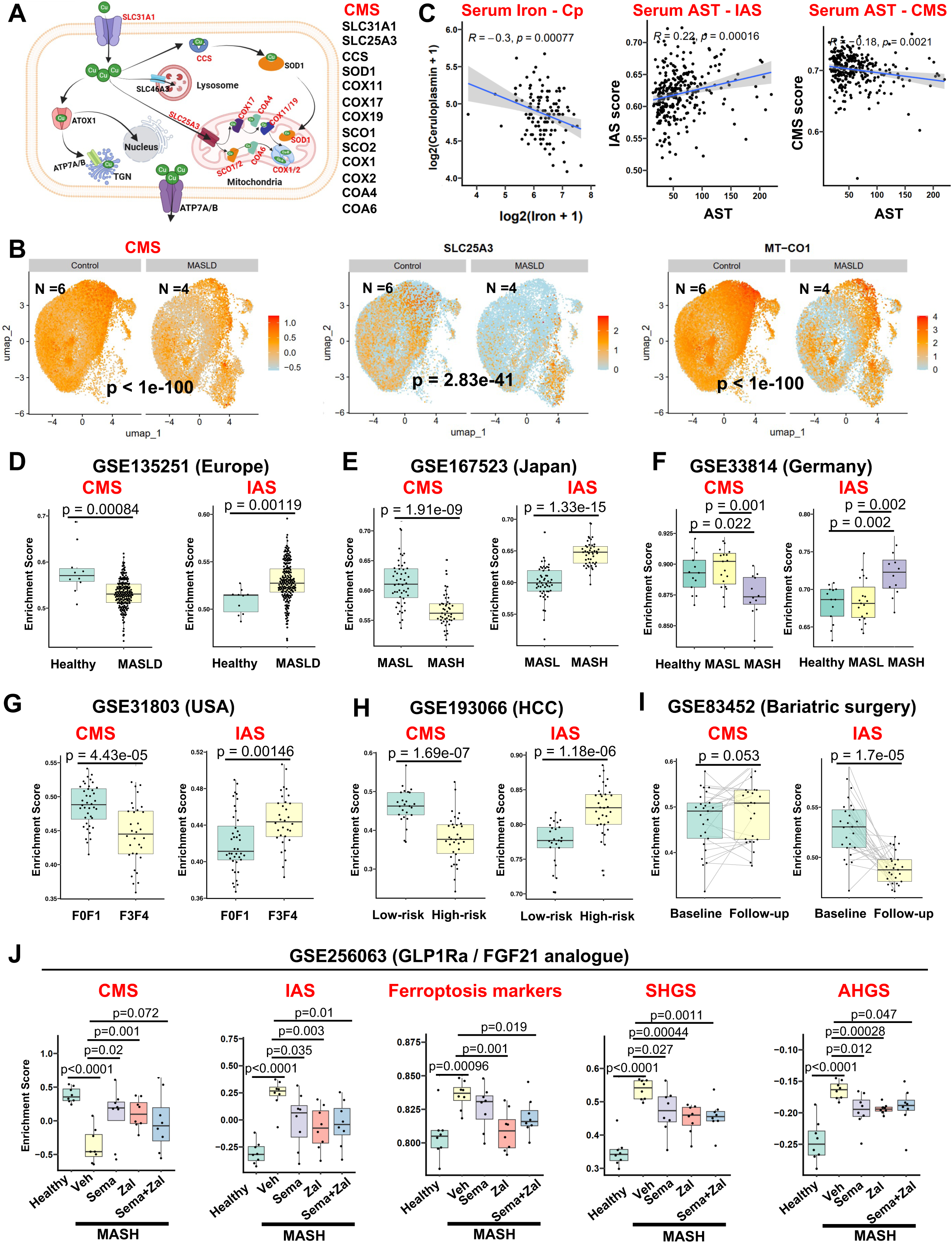
Copper-iron imbalance tracks MASLD progression, regression, and clinical outcomes. **(A)** Schematic illustrating the Cu-Mito Score (CMS) gene set representing mitochondrial copper sufficiency. **(B)** Single-cell RNA-seq visualization of CMS enrichment and expression of SLC25A3 and MT-CO1 in hepatocytes from control (n=6) and MASLD livers (n=4). **(C)** Correlations between serum iron, ceruloplasmin (Cp), serum AST, CMS, and the Iron Accumulation Score (IAS) in MASLD patients. **(D-G)** CMS and IAS enrichment across independent human liver transcriptomic cohorts, including European (GSE135251), Japanese (GSE167523), German (GSE33814) cohorts, and U.S. MASLD cohort (GSE31803). **(H)** CMS and IAS enrichment in MASLD patients stratified by hepatocellular carcinoma (HCC) risk (GSE193066). **(I)** Longitudinal CMS and IAS enrichment in paired liver biopsies obtained before and after bariatric surgery (GSE83452). **(J)** CMS, IAS, ferroptosis markers, and senescence-associated gene signature enrichment following treatment with a GLP-1 receptor agonist semaglutide or FGF21 analogue Zalfermin (GSE256063). Enrichment scores were calculated using GSVA; data are shown as box-and-whisker plots with individual data points; statistical comparisons were performed using Wilcoxon or *t* tests as appropriate, with *P* values shown.

At the systemic level, serum iron levels in MASLD patients negatively correlated with ceruloplasmin **(Fig. 5C)**, a liver-derived ferroxidase responsible for the majority of copper transport in circulation^23, 24^. CMS also inversely associated with serum AST, whereas IAS showed the opposite relationship **(Fig. 5C)**, supporting coordinated hepatic copper deficiency and iron excess during liver injury. Across multiple independent human liver cohorts spanning diverse geographic regions and disease stages, CMS progressively declined while IAS increased with MASLD severity, including transitions from healthy liver to steatosis, from MASLD to MASH, and from mild (F0-F1) to advanced fibrosis (F3-F4) **(Figs. 5D-G)**. Moreover, CMS inversely associated with key histologic features of disease activity, including steatosis grade, hepatocellular ballooning, portal inflammation, and lobular inflammation, whereas IAS increased across these same parameters **(Supplementary Fig. 7A)**. Copper-iron imbalance further tracked clinical outcomes and therapeutic response. In a MASLD-associated HCC cohort, patients classified as high risk exhibited significantly lower CMS and higher IAS than low-risk individuals, implicating mitochondrial copper deficiency and iron overload occur during malignant progression **(Fig. 5H)**. In a longitudinal bariatric surgery cohort, CMS increased while IAS decreased following weight loss and fibrosis regression, linking restoration of copper homeostasis and reduction of iron stress to disease improvement **(Fig. 5I)**. Similarly, in two independent studies, treatment with a GLP-1 receptor agonist semaglutide or FGF21 analogue Zalfermin increased CMS while suppressing IAS, ferroptosis, and senescence-associated signatures, indicating that effective metabolic therapy converges on copper-iron rebalancing **(Fig. 5J; Supplementary Fig. 7B)**.

Mechanistically, experimental labile iron overload models (either genetic or dietary) demonstrated that elevated IAS coincided with reduced CMS and increased ferroptosis and senescence signatures, and that these alterations were reversible by mitochondrial copper restoration with elesclomol **(Supplementary Fig. 8A, B; Fig. 4G)**, establishing copper-iron dyshomeostasis as a dynamic and therapeutically tractable axis. Together, these data demonstrate that mitochondrial copper deficiency and iron overload are tightly coupled features of MASLD progression, track histologic severity, predict treatment response and cancer risk, and can be reversed by metabolic or copper-targeted interventions.

### Copper-iron imbalance tracks MASLD-associated comorbidities, and restoring copper homeostasis suppresses kidney inflammation and fibrosis in MASH mice

Given the central roles of copper and iron in mitochondrial metabolism and cellular stress responses^22, 24^, we next examined whether copper-iron dysregulation extends beyond the liver to MASLD-associated extrahepatic comorbidities. We previously reported that senescence-associated gene signatures, including AHGS and SHGS, are enriched across multiple MASLD-related organ dysfunctions^25, 49, 51^, including heart and adipose tissue. Consistent with this, analysis of human heart failure cohorts revealed a marked reduction in CMS accompanied by increased IAS in both HFpEF and HFrEF compared with healthy controls, indicating coordinated mitochondrial copper insufficiency and iron overload in cardiac disease **(Fig. 6A)**. Similar reciprocal CMS-IAS patterns were observed in adipose tissue from individuals with metabolically unhealthy obesity compared with metabolically healthy states across two independent cohorts, linking copper-iron imbalance to adipose dysfunction associated with MASLD **(Figs. 6B, C)**. CMS and IAS were likewise dysregulated in other MASLD-associated degenerative conditions, including age-related macular degeneration and skeletal muscle sarcopenia, where reduced CMS coincided with increased inflammatory and senescence-related signaling **(Supplementary Figs. 9A, B)**.

**Fig. 6.**
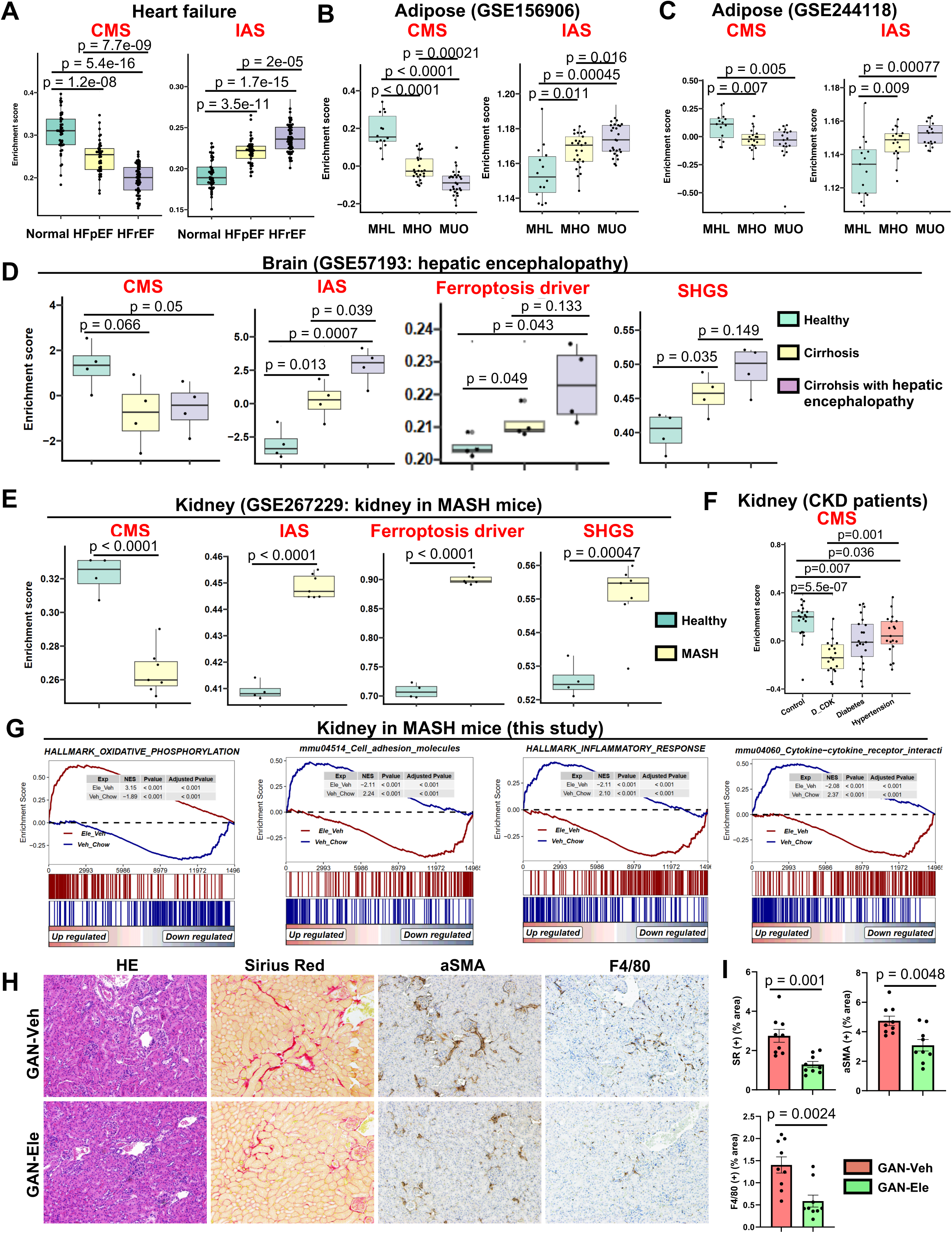
Copper-iron imbalance tracks MASLD-associated comorbidities, and restoring copper homeostasis suppresses kidney inflammation and fibrosis in MASH mice. **(A)** CMS and IAS enrichment scores in cardiac tissue from healthy controls, heart failure with preserved ejection fraction (HFpEF), and heart failure with reduced ejection fraction (HFrEF). **(B, C)** CMS and IAS enrichment in subcutaneous adipose tissue from metabolically healthy lean (MHL), metabolically healthy obese (MHO), and metabolically unhealthy obese (MUO) individuals in two independent cohorts (GSE156906 and GSE244118). **(D)** CMS, IAS, ferroptosis driver, and SHGS enrichment in brain tissue from controls, cirrhotic patients without hepatic encephalopathy, and cirrhotic patients with hepatic encephalopathy (GSE57193). **(E)** CMS, IAS, ferroptosis driver, and SHGS enrichment in kidney tissue from control and MASH mice (GSE267229). **(F)** CMS enrichment in kidney tissue from control subjects and patients with hypertension, diabetes, and diabetes-associated chronic kidney disease. **(G)** GSEA plots for oxidative phosphorylation, cell adhesion molecules, inflammatory response, and cytokine-cytokine receptor interaction pathways in kidneys from chow fed mice or GAN diet-fed mice treated with vehicle or elesclomol. **(H)** Representative kidney histology showing hematoxylin and eosin (H&E), Picrosirius Red, α-SMA, and F4/80 staining from GAN-fed mice treated with vehicle or elesclomol. **(I)** Quantification of renal fibrosis and inflammatory markers corresponding to panel H (data are shown as mean ± SEM). Enrichment scores were calculated using GSVA; data are shown as box-and-whisker plots with individual data points. Enrichment analyses in **A-G** were performed using permutation testing with Benjamini-Hochberg correction for multiple comparisons. Statistical significance in **I** was determined by Student t-test; *P* values are shown.

Copper-iron dysregulation was also evident in neurologic and renal complications linked to advanced liver disease. In cirrhotic patients with or without hepatic encephalopathy, CMS was reduced, whereas IAS, ferroptosis driver signatures, and senescence-associated hepatic gene signatures (including SHGS) were increased relative to healthy controls, suggesting heightened metal-driven cellular stress in neurologic dysfunction related to liver disease **(Fig. 6D)**. In parallel, kidney transcriptomes from MASH mice exhibited marked suppression of CMS together with elevated IAS, ferroptosis drivers, and SHGS, indicating that copper-iron imbalance accompanies renal injury in metabolic liver disease **(Fig. 6E)**. Consistent with these findings, CMS was also reduced in kidneys from patients with hypertension, diabetes, and chronic kidney disease, diseases that we previously showed were related to ferroptotic stress, accelerated aging and senescence^25, 49, 51^, underscoring strong translational relevance to human renal pathology **(Fig. 6F).**

To determine whether restoring copper homeostasis can mitigate renal injury, we analyzed kidneys from our GAN diet-fed MASH mice. GSEA revealed restoration of oxidative phosphorylation alongside suppression of fibrogenic, inflammatory, ferroptosis and cytokine/chemokine/IL6 signaling pathways in kidneys from elesclomol-treated mice compared with vehicle controls **(Fig. 6G; Supplementary Fig. 9C)**. Histologic analyses further demonstrated reduced renal fibrosis, macrophage infiltration, and inflammatory activation, as evidenced by decreased Sirius Red, α-SMA, and F4/80 staining **(Figs. 6H, I; Supplementary Fig. 9C)**. Collectively, these data indicate that mitochondrial copper repletion not only corrects hepatic copper-iron imbalance but also attenuates downstream renal inflammation and fibrosis *in vivo*. Together, these findings identify disrupted copper-iron homeostasis as a shared metabolic vulnerability across multiple organs and demonstrate that restoring mitochondrial copper homeostasis can suppress inflammation and fibrosis beyond the liver, with direct relevance to MASLD-associated systemic comorbidities.

## DISCUSSION

MASLD remains a leading driver of cirrhosis and HCC, yet the upstream determinants of disease progression remain incompletely understood. Disease progression continues in a substantial proportion of patients despite advances in our understanding of metabolic risk factors, underscoring gaps in how hepatocyte vulnerability to chronic metabolic stress is defined. Increasing evidence suggests that this vulnerability extends beyond lipid accumulation and insulin resistance and reflects impaired cellular resilience under sustained metabolic challenge^3^. Importantly, this impaired resilience is increasingly recognized as a systemic phenomenon that contributes to both hepatic injury and the extrahepatic comorbidities that drive adverse outcomes in MASLD^3^. One underexplored contributor is malnutrition and micronutrient imbalance, which are highly prevalent in chronic liver disease and independently predict morbidity and mortality^5, 6^. While most clinical and experimental studies have focused on macronutrients such as fat, calories, and protein, deficiencies in trace elements can exert disproportionate effects on mitochondrial function, redox balance, and cellular stress responses^5,6^. However, the mechanistic links between micronutrient dysregulation, hepatocyte fate, MASLD progression, and systemic metabolic vulnerability remain poorly defined.

Copper dysregulation represents a particularly compelling yet understudied facet of this vulnerability. Converging epidemiological and clinical evidence shows that copper in the Western diet has been decreasing at least since the 1930s, and half of the adult population consumes less than the amount recommended in the European Communities and the United Kingdom^14^. Hepatic copper levels are lower in MASLD than in healthy controls and even lower than levels in other chronic liver diseases^13^. Furthermore, copper deficiency associates with MASH severity, features of the metabolic syndrome, and adverse liver outcomes^13^. However, whether copper deficiency is a driver of disease or merely a consequence has remained unresolved. In a retrospective cohort of 183 patients with cirrhosis or portal hypertension, copper deficiency was associated with increased infection risk and a more than threefold higher risk of death, independent of age, sex, MELD-Na, and Karnofsky score^56^. These observations indicate that copper status is not simply a marker of disease severity but also an independent predictor of outcome. Preclinical studies corroborate the pathobiological significance of copper deficiency. In the ob/ob mouse model of MASLD, total hepatic metals appeared globally reduced; however, after lipid extraction, only copper remained selectively depleted^56^, suggesting a copper-specific deficit that may be masked by steatosis in bulk measurements. Dietary copper restriction in rodents is sufficient to induce hepatic steatosis and insulin resistance^13, 15^, while high-sucrose or high-fructose diets that promote steatohepatitis and liver fibrosis further worsen copper status and promote iron accumulation^16–18^. Notably, both copper deficiency and iron overload have been independently linked to multi-organ dysfunction including adipose tissue^57^, heart^58^, eye^59^, kidney^60^ and brain^61^. High-iron feeding caused systemic copper deficiency^62^, and copper supplementation reverses this dietary iron overload-induced multi-organ pathologies^63^, suggesting that trace metal imbalance may represent a shared pathogenic axis across organs rather than a liver-restricted abnormality. Further research is needed to tease apart mechanisms whereby liver injury *per se* impacts this process as liver is the major source of proteins that regulate systemic copper and iron homeostasis. In any case, the currently data indicate that copper dysregulation in MASLD is common, prognostically meaningful, at least partially reversible, and mechanistically linked to metabolic dysfunction, justifying focused investigation of hepatic copper homeostasis as an upstream and modifiable axis in organ damage related to systemic metabolic dysfunction.

A major advance from our work is the identification of mitochondrial copper insufficiency as a proximal cause of impaired cytochrome c oxidase integrity and bioenergetic failure in MASLD. Copper is indispensable for Complex IV assembly and function, yet its mitochondrial handling in metabolic liver disease has not been previously interrogated. By selectively restoring mitochondrial copper using the copper ionophore elesclomol, we demonstrate that copper deficiency is functionally reversible and that correcting this deficit restores respiratory competence, suppresses ferroptotic stress, and attenuates hepatocyte senescence and fibroinflammatory remodeling across complementary dietary models. These findings establish mitochondrial copper deficiency as a causal lesion, rather than a passive epiphenomenon, of liver injury.

Our data further reveal a feed-forward mechanism that amplifies copper deficiency once mitochondrial function is impaired. MASLD livers exhibit reduced expression of the mitochondrial copper transporter SLC25A3 and the copper-dependent Complex IV subunit MT-CO1, accompanied by altered signaling across the SLC25A3-SCO1-MTCO1-CTR1 axis. Prior studies have shown that defective COX assembly destabilizes CTR1 through SCO1-dependent signaling, thereby limiting cellular copper uptake^43–45^. Our findings support the operation of this mitochondria-to-plasma membrane feedback loop in MASLD, whereby mitochondrial copper insufficiency further restricts copper entry, amplifying bioenergetic failure and hepatocyte dysfunction. This self-reinforcing loop provides a mechanistic explanation for the persistence and progression of copper deficiency despite ongoing metabolic stress. Functional disruption of SLC25A3 in hepatocytes recapitulates these defects and exacerbates lipotoxic stress, directly linking impaired mitochondrial copper import to maladaptive hepatocyte fate.

An important implication of mitochondrial copper deficiency is its intersection with iron metabolism and ferroptosis. Copper is required for the ferroxidase activity of multicopper oxidases such as ceruloplasmin, which facilitate iron mobilization and prevent hepatic iron retention^23, 24^. Copper deficiency therefore creates a permissive environment for iron accumulation and lipid peroxidation, sensitizing hepatocytes to ferroptotic stress. By restoring mitochondrial copper, we suppress ferroptosis-associated programs and lipid peroxidation, providing a mechanistic bridge between copper deficiency, iron dysregulation, and ferroptotic injury, processes that have typically been considered independently in MASLD.

Beyond cell death pathways, our study identifies mitochondrial copper insufficiency as a determinant of hepatocyte identity and senescence. Senescent hepatocytes are increasingly recognized as active drivers of inflammation and fibrosis through secretion of senescence-associated mediators^4, 64^. We show that copper-deficient livers are transcriptionally depleted of mitochondrial bioenergetic and core hepatic metabolic programs and enriched for signatures of senescence, inflammation, and fibrosis. Restoring mitochondrial copper attenuates senescence markers and suppresses senescence-associated transcriptional programs in two preclinical models of MASH. Consistently, copper supplementation in palbociclib-induced senescent Huh7 cells reduced β-galactosidase positivity and suppressed canonical SASP factors. Together, these findings demonstrate that copper availability directly regulates hepatocyte senescence and SASP activity, independent of systemic metabolic effects, and place micronutrient status upstream of hepatocyte fate decisions and microenvironmental remodeling in MASLD.

The translational relevance of these findings is reinforced by our development of the CMS, a transcriptome-based metric of mitochondrial copper sufficiency. Across multiple independent human cohorts, lower CMS associates with iron overload, MASLD severity, diminished therapeutic response, and increased HCC risk. Importantly, CMS and IAS exhibit coordinated dysregulation across multiple extrahepatic tissues implicated in MASLD-associated comorbidities. These observations suggest that disrupted copper-iron homeostasis reflects a systemic metabolic vulnerability. Given the liver’s central role in regulating systemic copper and iron distribution through proteins such as ceruloplasmin, and its broader function as a master regulator of whole-body metabolism, hepatic copper–iron imbalance may act as a nodal amplifier of multi-organ dysfunction in MASLD. Accordingly, CMS and IAS may have utility not only as liver-focused biomarkers but also as tools to assess multi-organ risk and therapeutic response. Correcting copper-iron imbalance may therefore represent an opportunity to attenuate both liver disease progression and the systemic comorbidities that critically shape MASLD outcomes.

Several limitations warrant consideration. First, although SLC25A3 is required for mitochondrial copper delivery in mammalian cells, it is best known as a mitochondrial phosphate carrier in yeast^65, 66^, raising the possibility that the observed bioenergetic defects could reflect impaired phosphate transport rather than copper imbalance. Multiple lines of evidence argue against this interpretation. *Slc25a3* expression declines in MASLD hepatocytes, and selective restoration of mitochondrial copper using elesclomol rescues mitochondrial bioenergetics and attenuates disease phenotypes without manipulating phosphate availability. In addition, prior studies demonstrate that cytochrome c oxidase defects resulting from SLC25A3 loss in mammalian systems are rescued by copper, not phosphate in mammalian cells^40^. Nevertheless, future studies that independently manipulate mitochondrial copper and phosphate availability *in vivo* will be valuable to further refine causal attribution.

Safety considerations are also important. At high concentrations, elesclomol-copper complexes can induce cuproptosis in cancer contexts^36^, raising theoretical concerns regarding therapeutic application. Several safeguards mitigate this risk in the present study. The concentrations of elesclomol used to rescue hepatocyte lipotoxicity are low and well below those reported to induce oxidative stress or cuproptosis. Elesclomol has demonstrated efficacy and tolerability in genetic copper-deficiency models^33, 35, 67^, including clinical benefit in children with Menkes disease^68^. Moreover, in both MASLD dietary models examined here, elesclomol reduced liver injury rather than increasing cell death, and we did not observe induction of cuproptosis-associated transcriptomic programs. Nonetheless, careful dose optimization and longitudinal monitoring of cuproptosis markers will remain important considerations as copper-targeted strategies advance toward clinical translation.

In summary, our study identifies mitochondrial copper deficiency as a central and previously underappreciated driver of MASLD pathogenesis. By linking micronutrient imbalance to mitochondrial dysfunction, iron-dependent ferroptotic stress, hepatocyte senescence, and fibroinflammatory remodeling, this work establishes copper-centered mitochondrial regulation as a unifying mechanistic framework for disease progression. Targeting mitochondrial copper homeostasis may therefore represent a novel therapeutic strategy to restore hepatocyte bioenergetic resilience, disrupt maladaptive cell fate programs, and attenuate progression of MASLD and its downstream complications and multiorgan dysfunctions.

## METHODS and MATERIALS

### Mice and husbandry

Male C57BL/6J mice were obtained from The Jackson Laboratory (Bar Harbor, ME) and housed in a specific pathogen-free barrier facility under a 12 h:12 h light-dark cycle with ad libitum access to food and water. Unless otherwise specified, mice were 8-12 weeks of age at the start of dietary interventions. All animal procedures were approved by the Duke University Institutional Animal Care and Use Committee (IACUC) and were conducted in accordance with NIH guidelines for humane animal care.

### Dietary models of MASLD and elesclomol treatment

To model MASLD across distinct stages and mechanisms, male C57BL/6J mice were subjected to complementary dietary regimens. For advanced steatohepatitis, mice were fed a GAN diet (D09100310; Research Diets) for 34 weeks, while control mice received standard chow. To evaluate therapeutic efficacy in established disease, GAN-fed mice were randomized to receive vehicle (10% Cavitron™ W7 HP7 Pharma cyclodextrin) or the copper ionophore elesclomol (10 mg/kg, oral gavage, three times per week) during the final 5 weeks of dietary exposure. To assess generalizability, an independent cohort of mice was fed a choline-deficient, L-amino acid-defined high-fat diet (CDAHFD; A06071302, Research Diets) for 10 weeks. CDAHFD-fed mice similarly received vehicle or elesclomol during the final 5 weeks of diet administration using the same dosing regimen. At study completion, blood was collected, and liver tissues were either fixed in buffered formalin for histologic and immunohistochemical analyses or snap-frozen in liquid nitrogen and stored at -80 °C for transcriptomic, biochemical, and ICP-MS based metallomic analyses.

### Mitochondrial Isolation

For assessment of the effects of elesclomol on hepatic and mitochondrial copper levels, mice maintained on a standard chow diet were randomly assigned to receive either elesclomol or vehicle (n = 4 mice per group). Elesclomol was administered by oral gavage at 10 mg/kg once every other day for one week (4 doses in total), while control mice received vehicle alone (10% Cavitron™ W7 HP7 Pharma cyclodextrin). At the end of the treatment period, livers were harvested for subcellular fractionation by differential centrifugation under cold conditions^69^.

Briefly, right and caudate liver lobes were excised, finely minced, and homogenized on ice in isotonic isolation buffer (pH 7.4) containing mannitol, sucrose, HEPES, EDTA, EGTA, and fatty acid–free bovine serum albumin. Homogenization was performed using a motor-driven Teflon pestle until uniform disruption was achieved. The homogenate was first cleared of unbroken cells and nuclei by low-speed centrifugation (2,500 × g for 10 min at 4 °C). The resulting supernatant was then centrifuged at high speed (20,000 × g for 10 min at 4 °C) to pellet crude mitochondria, while the supernatant was retained as the cytosolic fraction. Mitochondrial pellets were gently washed once with isolation buffer to minimize cytosolic contamination, followed by two washes with ice-cold PBS to reduce residual buffer components, and immediately snap-frozen in liquid nitrogen. Both whole liver tissue and isolated mitochondrial fractions were stored at −80 °C.

### ICP-MS analysis

Elemental copper concentrations were quantified in whole liver tissue and isolated mitochondrial fractions by inductively coupled plasma mass spectrometry (ICP-MS). Frozen samples were weighed and digested in trace metal–grade nitric acid (HNO₃; Fisher Chemical) at elevated temperature until complete dissolution was achieved. Digested samples were diluted in ultrapure water to appropriate final acid concentrations for ICP-MS analysis. ICP-MS measurements were performed using an Agilent ICP-MS system (Agilent Technologies, Santa Clara, CA) operated under standard conditions with collision/reaction cell technology to minimize polyatomic interferences. Calibration curves were generated using multi-element standard solutions (Inorganic Ventures or equivalent) spanning the expected concentration range. Internal standards were included to correct for instrumental drift and matrix effects, and blanks were run in parallel to control for background contamination. For whole liver samples, copper concentrations were normalized to tissue dried weight and expressed as micrograms of metal per gram of tissue. For mitochondrial fractions, elemental content was normalized to mitochondrial wet weight and expressed as nanograms of metal per milligram of mitochondrial pellets.

### Cell culture and *in vitro* assays

AML12 mouse hepatocytes (ATCC) were maintained in DMEM/F12 medium supplemented with 10% fetal bovine serum, 1% penicillin/streptomycin, insulin-transferrin-selenium, and dexamethasone under standard culture conditions (37 °C, 5% CO₂). To model hepatocyte lipotoxicity, cells were treated with a mixture of oleic acid and palmitic acid (OPA; 200 μM oleate and 100 μM palmitate) complexed to fatty acid-free bovine serum albumin. Control cells received vehicle (BSA) alone. To assess the effects of mitochondrial copper restoration, cells were co-treated with elesclomol (0.25 nM) during OPA exposure. For mechanistic studies, *Slc25a3* expression was silenced using siRNA transfection (10 μM; Dharmacon™ ON-TARGETplus siRNA) prior to OPA treatment, with non-targeting siRNA serving as control. Cells were maintained under lipotoxic conditions for up to 6 days, with media refreshed every 3 days. At the end of treatment, cells were harvested for immunoblotting, lipid staining, and transcriptomic analyses. Knockdown efficiency was confirmed by immunoblotting and quantitative PCR. Mitochondrial integrity and respiratory chain protein abundance were evaluated by immunoblotting for representative subunits across Complexes I-V. Lipid accumulation was assessed by Oil Red O staining and images were acquired using Leica Microsystems.

To determine whether copper availability directly regulates hepatocyte senescence independent of systemic effects, senescence was induced in human Huh7 hepatocytes (a gift from Dr. Charles M. Rice, Rockefeller University) using the CDK4/6 inhibitor palbociclib (1 μM; MedChemExpress) for 3 days. Where indicated, cells were subsequently treated with copper sulfate (CuSO₄; 5 μM). Senescent cells were measured by Senescence-associated β-galactosidase staining (#9860, Cell Signaling), and expression of canonical senescence-associated secretory phenotype (SASP) factors, including *CXCL1*, *CXCL2*, *IL8*, and *TNF*α, was assessed by quantitative PCR.

### Quantitative real-time PCR (qRT-PCR)

Total RNA was isolated from snap-frozen liver, kidney tissue or cultured cells using TRIzol reagent (Thermo Fisher Scientific, Waltham, MA) according to the manufacturer’s instructions. RNA concentration and purity were assessed spectrophotometrically. Complementary DNA (cDNA) was synthesized from equal amounts of total RNA using SuperScript™ II Reverse Transcriptase (Life Technologies, Carlsbad, CA). Quantitative real-time PCR was performed using SYBR™ Green Supermix (Life Technologies) on a QuantStudio™ 6 Real-Time PCR System (Thermo Fisher Scientific). Gene-specific primer sequences are provided in Supplementary Table 1. Transcript levels were normalized to the ribosomal housekeeping gene *S9*, and relative gene expression was calculated using the comparative Ct (2^−ΔΔCt^) method.

### Immunoblot analysis

Protein lysates were prepared from whole liver tissue or cultured cells using RIPA buffer supplemented with protease inhibitors (Sigma-Aldrich, St. Louis, MO). Protein concentrations were determined prior to electrophoresis, and equal amounts of protein were resolved on 4-20% gradient SDS–PAGE gels (Criterion™ TGX™, Bio-Rad, Hercules, CA). Proteins were transferred onto PVDF membranes and incubated with primary antibodies listed in Supplementary Table 2. All antibodies were selected based on prior validation for species and application, as documented by manufacturers and were further optimized and titrated in-house using appropriate positive or negative biological controls. Membranes were incubated with horseradish peroxidase-conjugated secondary antibodies, and immunoreactive bands were visualized using Image Studio™ Lite software (version 5.2; LI-COR Biosciences, Lincoln, NE).

### Histology and immunohistochemistry (IHC)

Liver and kidney tissues were fixed in phosphate-buffered formalin, paraffin-embedded, and sectioned for histopathologic analysis. Hematoxylin and eosin (H&E) staining was performed to assess overall liver architecture and steatohepatitis. Fibrosis was evaluated using Picrosirius Red staining (Sigma-Aldrich, 365548). For immunohistochemistry, paraffin sections were deparaffinized, rehydrated, and incubated with 3% hydrogen peroxide for 10 minutes to quench endogenous peroxidase activity. Antigen retrieval was performed by heating sections in 10 mmol/L sodium citrate buffer (pH 6.0) for 10 minutes. Sections were blocked with Dako protein block solution (Agilent Technologies, Santa Clara, CA) for 1 hour and incubated overnight at 4 °C with primary antibodies listed in Supplementary Table 2. Polymer-based horseradish peroxidase secondary antibodies were applied for 1 hour at room temperature, followed by detection using the Dako 3,3′-diaminobenzidine (DAB) substrate chromogen system. Images were acquired using a Zeiss LSM710 imaging platforms and quantified using ImageJ software.

### Bulk RNA-seq library preparation and preprocessing

Total RNA was isolated from liver, kidney tissue or cultured cells using TRIzol reagent (Thermo Fisher Scientific, Waltham, MA). Bulk RNA-seq libraries were prepared and sequenced by Novogene on an Illumina NovaSeq platform using 150-bp paired-end reads. Raw sequencing reads were subjected to quality control, adaptor trimming, and removal of low-quality bases prior to alignment using *fastp*^70^. Cleaned reads were aligned to the mouse reference genome (mm39) using STAR (v2.7.5). Gene-level read counts were quantified using *featureCounts* based on standard gene annotations^71^.

### Differential expression and pathway enrichment analyses

Differential gene expression was analyzed using R DESeq2 package^72^. Lowly expressed genes were filtered prior to model fitting, and the significance of differential expression was processed with multiple-testing correction using the Benjamini-Hochberg method. Gene set enrichment analyses (GSEA) were performed using R *clusterProfiler* (v61) on preranked gene lists derived from DESeq2^73^. Curated pathway collections, including Reactome, Hallmark, and KEGG gene sets, were obtained from the Molecular Signatures Database (MSigDB) using the R *msigdbr* package.

### Development and application of the Cu-Mito Score (CMS)

#### CMS gene signature construction

To evaluate mitochondrial copper sufficiency, we developed a Cu-Mito Score (CMS) gene signature, composed of genes regulating cellular copper uptake, mitochondrial copper transport, cytochrome c oxidase (COX) assembly, and copper-dependent redox homeostasis^54, 74^. Candidate genes were curated based on their known roles in copper handling and mitochondrial bioenergetics function. The final CMS consisted of 13 genes, including copper transporters (e.g., *SLC31A1*, *SLC25A3*), COX subunits and assembly factors (*COX1*, *COX2*, *COX17*, *COX19*, *SCO1*, *SCO2*), and copper chaperones (*CCS*, *SOD1*), capturing mitochondrial copper availability and functional utilization^54, 74^ (Fig. 5A).

In parallel, iron burden was assessed using a previously validated Iron Accumulation Score (IAS), derived from a 128-gene signature enriched for iron uptake, storage, and iron-responsive stress programs^55^ (Fig. 5B). CMS and IAS were analyzed as complementary but biologically distinct axes reflecting mitochondrial copper sufficiency and iron overload, respectively.

### CMS and IAS enrichment analysis in human MASLD cohorts

For bulk transcriptome datasets, CMS and IAS enrichment scores were calculated using Gene Set Variation Analysis (GSVA) using R *GSVA* package. For microarray datasets, raw data were downloaded using the R *GEOquery package (cite)* and normalized using GCRMA or RMA method. For bulk RNA-seq datasets, raw gene count matrix was normalized using DESeq2 prior to GSVA.

CMS and IAS enrichment was evaluated across multiple independent human MASLD cohorts spanning disease severity, therapeutic response, and cancer risk:

1. **Duke MASLD cohort** (GSE213623; n = 368). This cohort included individuals with obesity without MASLD (n = 69) and individuals with obesity and biopsy-proven MASLD (n = 299). Liver tissue, blood samples, and clinical data were collected through the Duke University Health System (DUHS) MASLD Clinical Database and Biorepository. Bulk liver RNA-seq data were used for CMS and IAS enrichment analyses, which were correlated with clinical parameters including serum albumin, AST, and Fib-4 score.
2. **German MASLD cohort** (GSE33814; n = 44). Liver transcriptomes from normal liver (n = 13), simple steatosis (n = 18), and steatohepatitis (n = 12) were generated using Affymetrix HG-U133 Plus 2.0 microarrays. Raw CEL files were downloaded using GEOquery and normalized using the gcrma method prior to CMS and IAS enrichment analysis.
3. **Japanese MASLD cohort** (GSE167523; n = 98). This retrospective cohort included patients with simple steatosis (n = 51) or non-alcoholic steatohepatitis (n = 47). Bulk RNA-seq count matrices were obtained from NCBI GEO, normalized using DESeq2, and analyzed for CMS and IAS enrichment.
4. **European MASLD cohort** (GSE135251; n = 216). Bulk RNA-seq was performed on liver biopsies from 206 MASLD patients and 10 healthy controls across France, Germany, Italy, and the United Kingdom. Fibrosis severity ranged from F0 to F4. Raw counts were normalized using DESeq2 prior to CMS and IAS deconvolution.
5. **MASLD-HCC primary risk cohort** (GSE193066; n = 106). This cohort comprised non-cirrhotic, HCC-naïve MASLD patients who underwent diagnostic liver biopsy at Hiroshima University and were followed prospectively for a median of 8.9 years. CMS and IAS enrichment were correlated with HCC risk, defined using an etiology-agnostic prognostic liver signature (PLS).
6. **Bariatric surgery follow-up cohort** (GSE83452; n = 25). Paired liver biopsies obtained at baseline and one year following bariatric surgery were analyzed using Affymetrix HG-U133 Plus 2.0 arrays. Raw CEL files were normalized using R *gcrma* package, enabling longitudinal assessment of CMS and IAS changes in response to metabolic improvement.

### CMS and IAS analysis in extrahepatic tissues

To determine whether mitochondrial copper insufficiency and iron overload represent broader features of chronic metabolic or degenerative disease beyond the liver, CMS and IAS enrichment was evaluated across multiple transcriptomic datasets from extrahepatic human tissues. These cohorts encompassed adipose tissue, heart, kidney, brain, eye, and skeletal muscle, each representing clinically relevant states of metabolic dysfunction, organ failure, or fibrosis. Across these extrahepatic tissues, CMS and IAS were analyzed to assess whether mitochondrial copper insufficiency and iron overload represent shared transcriptional programs associated with metabolic stress, organ dysfunction, or degenerative disease states.

1. **Subcutaneous adipose tissue cohort 1** (GSE244118; n = 53). Bulk RNA-seq was performed on subcutaneous abdominal adipose tissue from individuals categorized as metabolically healthy lean (MHL; n = 15), metabolically healthy obese (MHO; n = 19), and metabolically unhealthy obese (MUO; n = 19). MHO status was defined by normal fasting glucose, glucose tolerance, HbA1c, intrahepatic triglyceride content, serum triglycerides, and whole-body insulin sensitivity, whereas MUO was defined by prediabetes and hepatic steatosis. Raw counts were normalized using DESeq2, followed by CMS and IAS enrichment analysis using GSVA.
2. **Subcutaneous adipose tissue cohort 2** (GSE156906; n = 56). Bulk RNA-seq data were analyzed from subcutaneous abdominal adipose tissue of MHL (n = 14), MHO (n = 25), and MUO (n = 17) individuals. DESeq2 normalization and GSVA were applied to quantify CMS and IAS enrichment across metabolic states.
3. **Heart failure cohort** (Zenodo; n = 145). Gene count data were obtained from left or right ventricular septal endomyocardial biopsies from patients with heart failure with preserved ejection fraction (HFpEF; n = 41), heart failure with reduced ejection fraction (HFrEF; n = 59), and healthy controls (n = 45). Raw counts were normalized using DESeq2, and CMS and IAS enrichment scores were calculated using GSVA.
4. **Kidney cohort** (E-MTAB-2502; n = 79). Microarray data were obtained from human kidney tubule samples classified as control (n = 20), diabetes–chronic kidney disease (D-CKD; n = 19), diabetes without CKD (n = 21), and hypertension (n = 19). Data were generated using the Affymetrix HG-U133A_2 array. For genes represented by multiple probes, mean probe expression was used. CMS and IAS enrichment was performed using GSVA.
5. **Brain cohort (**GSE57193; n=12). Gene expression of post mortem human fusiform gyrus samples were profiled by Agilent Whole Human Genome Microarrays (GPL14550) from healthy controls (n=4), cirrhosis without hepatic encephalopathy (n=4), and cirrhosis with hepatic encephalopathy (n=4). Gene expression data were processed using R *limma* package and transformed with quantile-normalized method. For probes were mapped to genes, they were collapsed by mean when multiple probes mapped to the same gene. CMS and IAS enrichment was computed using GSVA.
6. **Age-related macular degeneration** (GSE115828; n=523). Retinal tissue transcriptome data were obtained from 523 donors spanning controls and distinct stages of age-related macular degeneration (AMD) patients. Following original publication^75^, the disease status is defined using Minnesota Grading System (MGS). Patients with MGS level of 1 were set as control, and were grouped as AMD if MGS >1. DESeq2 was used to normalize gene counts, and GSVA for computing CMS and IAS score.
7. **Sarcopenia** (GSE226151; n=60). Bulk RNA-seq data were obtained from human skeletal muscle samples across three groups: healthy (n = 20), pre-sarcopenia (n = 20), and sarcopenia (n = 20). Short reads were sequenced on an Illumina HiSeq 2500 platform, and the gene-level read count matrix was downloaded from NCBI GEO. Counts were normalized by DESeq2, and CMS and IAS enrichment scores were calculated using GSVA.

### Single-nucleus RNA-seq processing and cell-type annotation

Single-nucleus RNA-seq data were processed from raw sequencing output to generate nucleus-by-gene count matrices using standard preprocessing pipelines. Downstream analyses were performed using Seurat (v5.1.0)^76^. Nuclei were filtered based on quality-control metrics, including the number of detected genes, UMI counts, and mitochondrial gene content. Data normalization, scaling, dimensionality reduction, and batch integration were performed using Seurat’s standard workflows. Cell clusters were visualized by UMAP and annotated based on canonical marker genes.

### Hepatocyte-focused analyses and Cu-Mito Score stratification

Hepatocytes were subset for downstream analyses. To quantify mitochondrial copper sufficiency at single-cell resolution, the CMS score was calculated using AUCell^77^. For each hepatocyte, AUCell calculated enrichment scores based on expression ranking of CMS genes, and hepatocytes were stratified into CMS-high and CMS-low populations using an adjusted AUC threshold. Differential expression and pathway enrichment analyses were then performed comparing CMS-low versus CMS-high hepatocytes.

Gene set enrichment analyses in hepatocyte subsets were conducted using clusterProfiler with Reactome, Hallmark, and KEGG gene sets, with emphasis on mitochondrial metabolic programs, core hepatocyte functions, epithelial–mesenchymal transition, ferroptosis, inflammation, fibrosis, and senescence-associated pathways.

## Statistical analysis

All experimental data were analyzed using GraphPad Prism (v10.2.1; GraphPad Software) or Microsoft Excel (Microsoft Office 2021). Data are presented as mean ± SEM unless otherwise indicated. Comparisons between two groups were performed using two-tailed Student’s *t* tests. Comparisons among multiple groups were assessed by one-way analysis of variance (ANOVA) followed by Tukey’s post hoc multiple-comparison test. A *p* value < 0.05 was considered statistically significant. For transcriptomic analyses, data visualization was performed in R using *ggplot2*. Differences in gene-set enrichment scores between groups were evaluated using either the Wilcoxon rank-sum test or Student’s *t* test, as appropriate, implemented via the *ggpubr* package.

## Data availability

All data generated in this study are included in this article and its supplementary materials. Newly generated bulk RNA-seq and single-nucleus RNA-seq raw sequencing files and processed count matrices will be deposited in the NCBI Gene Expression Omnibus (GEO) upon manuscript acceptance.

## Code Availability

The algorithms used for the analysis in this study are all publicly available. No new algorithm or software was developed from this study.

## Acknowledgements

This work was supported by the NIDDK K01DK135793-01A1 to K.D., P20GM113226(Sub-Project ID 6813) and R21CA290420 to M.S., and National Institutes of Health grants R01 AA010154, R01 DK077794, R56 DK134334 and Sponsored Research Study Agreement 337521 supported by Boehringer Ingelheim Pharmaceuticals, Inc, awarded to A.M.D. The authors are grateful to the patients who donated liver tissue for analysis, to Drs. Manal Abdelmalek and Cynthia Guy and the clinical staff and coordinators who created and maintain the Duke NAFLD Biorepository, Emily Hocke, Stephanie Arvai, and Karen Abramson from the Duke Molecular Genetics Core for single nuclei RMA sequencing of control and MASLD liver tissues, Dr. Steven Pullen’s team at Boehringer Ingelheim Pharmaceuticals, Inc. for their assistance in the bioinformatics analysis.

## Author Contributions

K.D., L.W. and N.R. conceived of the experiments. N.R, K.D. and M.S. performed experiments. N.R, L.W., R.K.D, D.S.U, S.H.O, D.C.K., A.M.D and K.D. analyzed data. K.D., L.W. and A.M.D. wrote the manuscript. K.D. and A.M.D. secured funding for the study. Everyone reviewed and approved the manuscript. Correspondence to A.M.D. or K.D.

## Declaration of Generative AI and AI-assisted technologies in the writing process

During the preparation of this work, the authors used ChatGPT (OpenAI) to correct grammatical errors and enhance clarity and readability. After using this tool, the authors reviewed and edited the content as needed and take full responsibility for the content of the publication.

---

**Supplementary Fig. 1.**
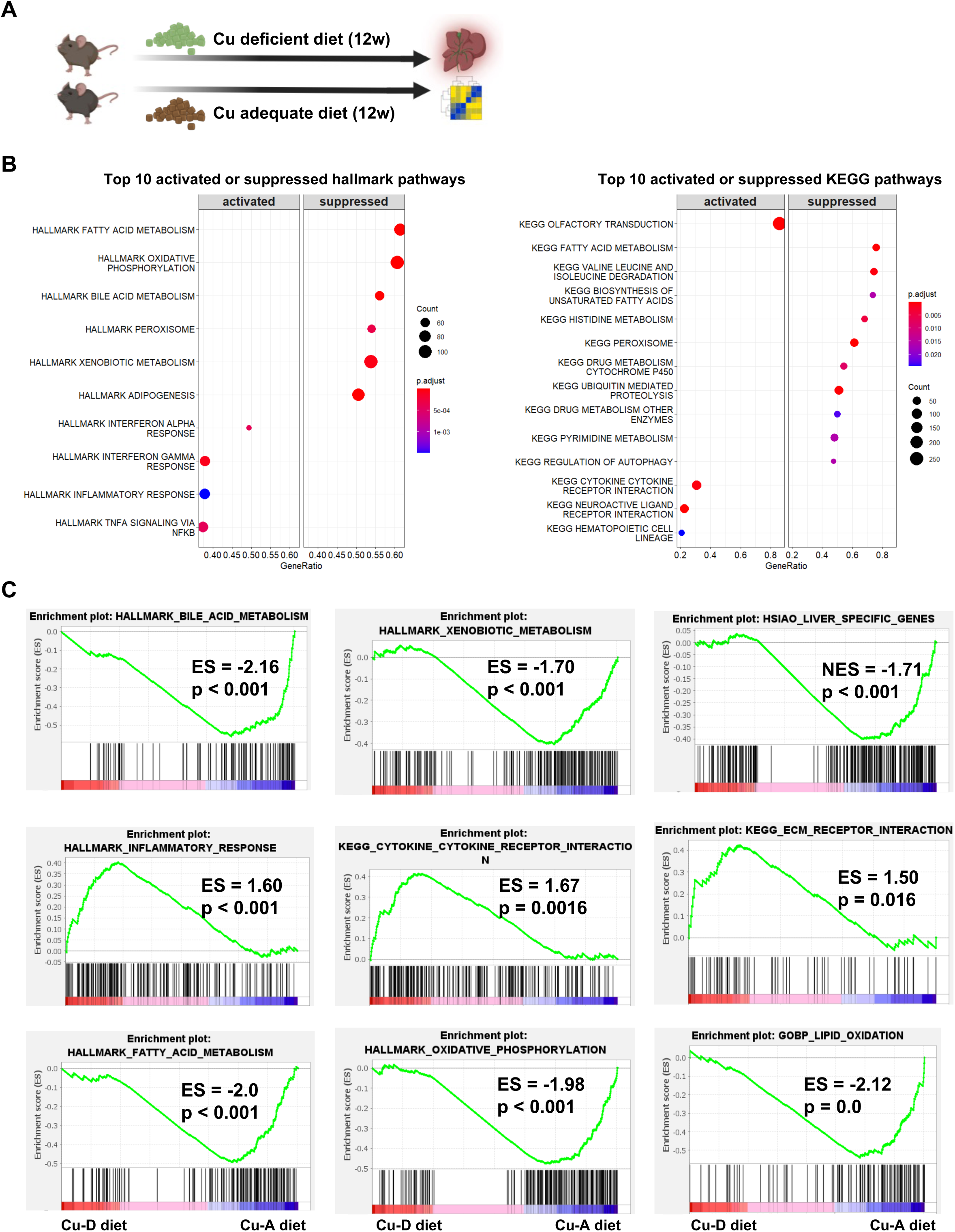
Dietary copper deficiency induces MASLD-like transcriptomic changes. **(A)** Experimental design. Adult rodents were fed a copper-deficient AIN-76A diet (Cu-D; <0.3 mg/kg copper) or a copper-adequate control diet (Cu-A; 125 mg/kg copper) for 12 weeks (GSE58875). **(B)** Gene set enrichment analysis (GSEA) showing the top 10 activated and suppressed Hallmark (left) and KEGG (right) pathways in liver transcriptomes from Cu-D versus Cu-A animals. **(C)** Representative GSEA enrichment plots for selected Hallmark and KEGG pathways. P values were calculated using permutation testing and adjusted for multiple comparisons using the Benjamini-Hochberg method.

**Supplementary Fig. 2.**
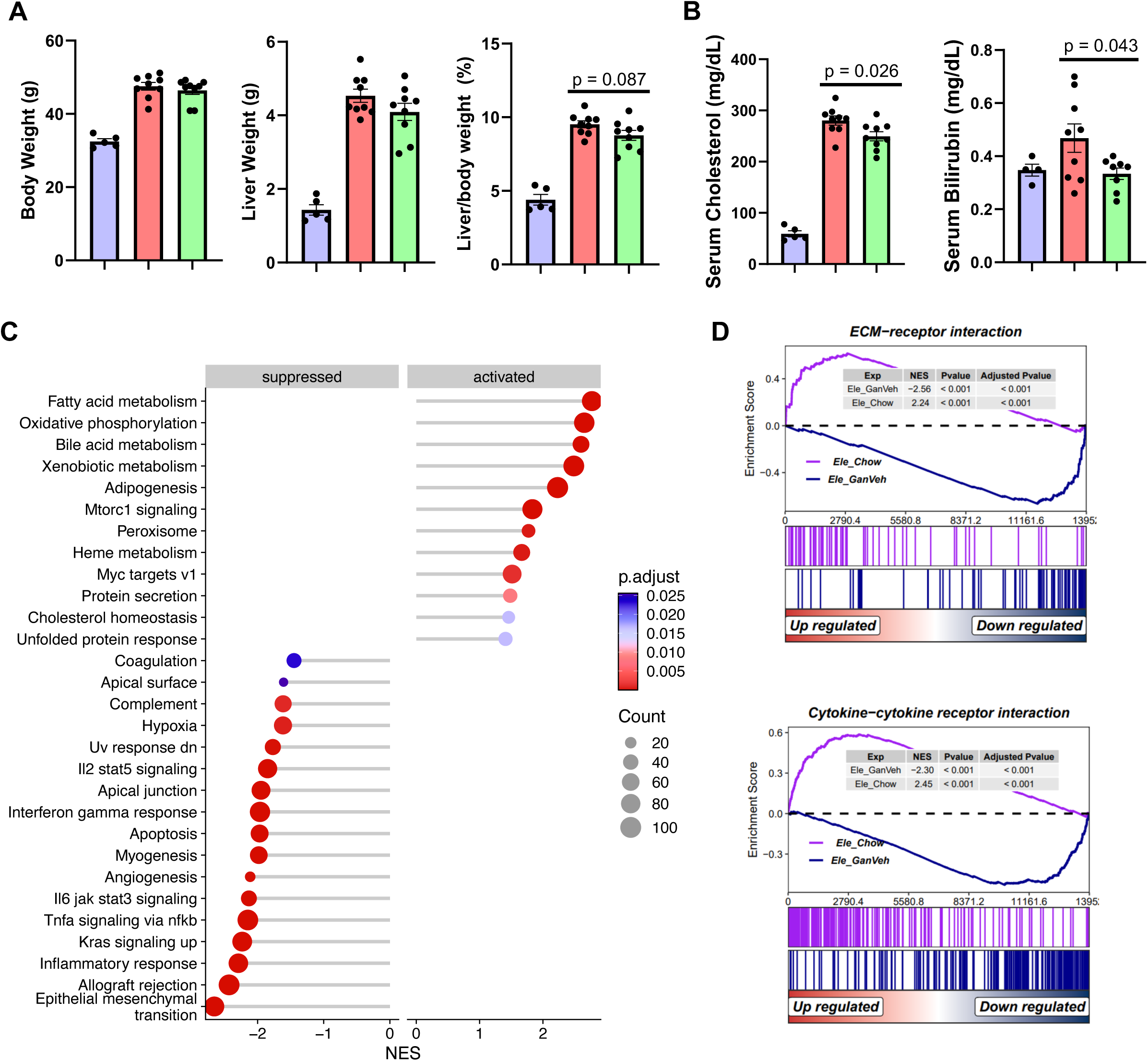
Mitochondrial copper ionophore elesclomol ameliorates MASLD in GAN diet–fed mice. **(A)** Body weight, liver weight, and liver-to-body weight ratio in chow-fed mice and GAN diet-fed mice treated with vehicle or elesclomol during the final 5 weeks of diet exposure. **(B)** Serum total cholesterol and bilirubin levels across experimental groups. Data are shown as mean ± SEM. **(C)** Gene set enrichment analysis (GSEA) showing normalized enrichment scores (NES) for activated and suppressed Hallmark pathways in GAN-Ele versus GAN-Veh livers. **(D)** GSEA enrichment plots for KEGG ECM-receptor interaction and cytokine-cytokine receptor interaction pathways comparing GAN-Ele with GAN-Veh. Statistical significance in A-B was assessed by one-way ANOVA with post hoc testing as indicated. GSEA *P* values were calculated using permutation testing and adjusted for multiple comparisons using the Benjamini-Hochberg method.

**Supplementary Fig. 3.**
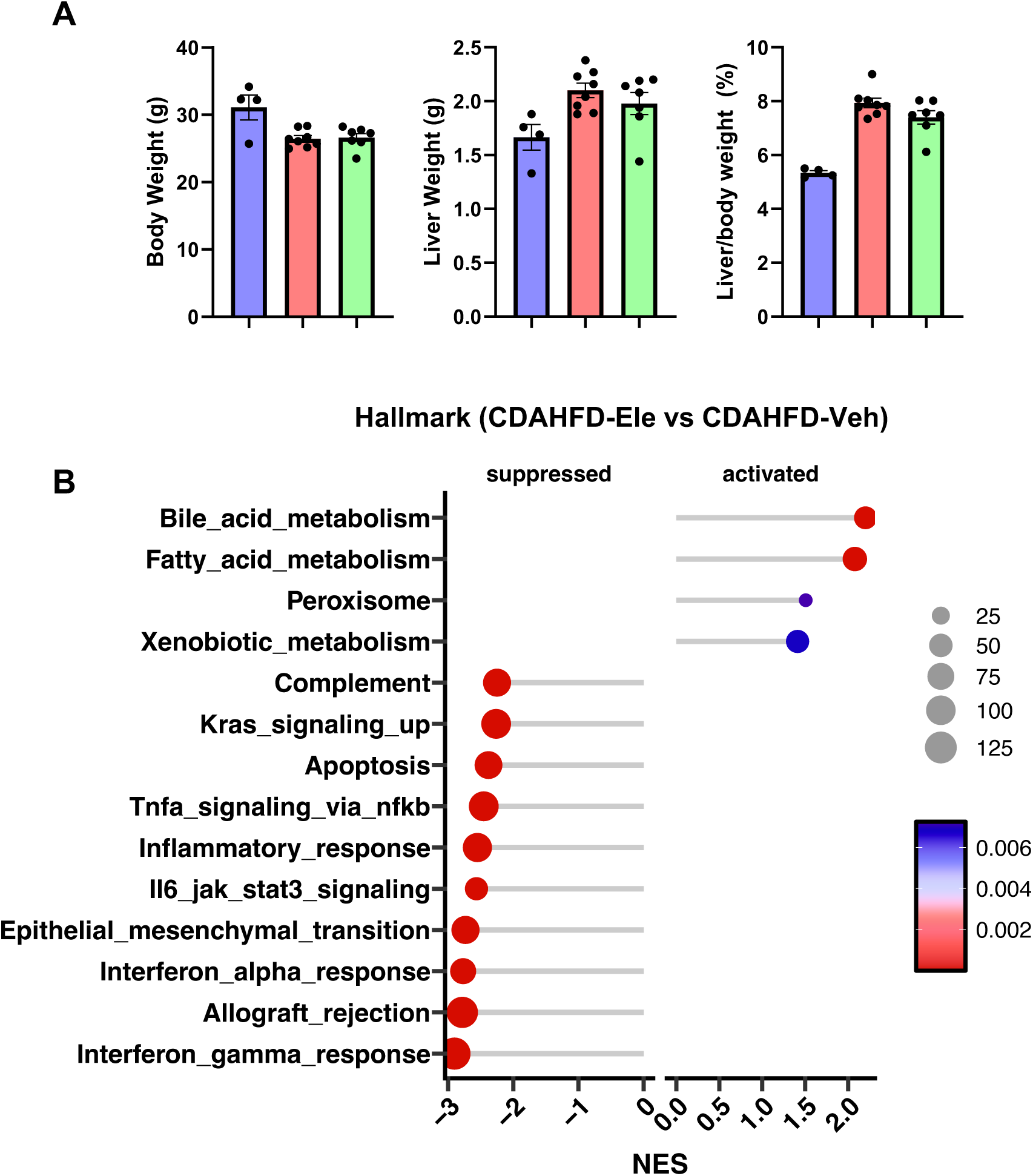
Elesclomol ameliorates MASLD in CDAHFD-fed mice. **(A)** Body weight, liver weight, and liver-to-body weight ratio in chow-fed mice and CDAHFD-fed mice treated with vehicle or elesclomol (10 mg/kg, oral gavage, three times per week during the final 5 weeks of diet exposure). **(B)** Gene set enrichment analysis (GSEA) of Hallmark pathways comparing liver transcriptomes from CDAHFD-Ele versus CDAHFD-Veh mice, showing the top activated and suppressed pathways. Data in A are shown as mean ± SEM. GSEA *P* values were calculated using permutation testing and adjusted for multiple comparisons using the Benjamini-Hochberg method.

**Supplementary Fig. 4.**
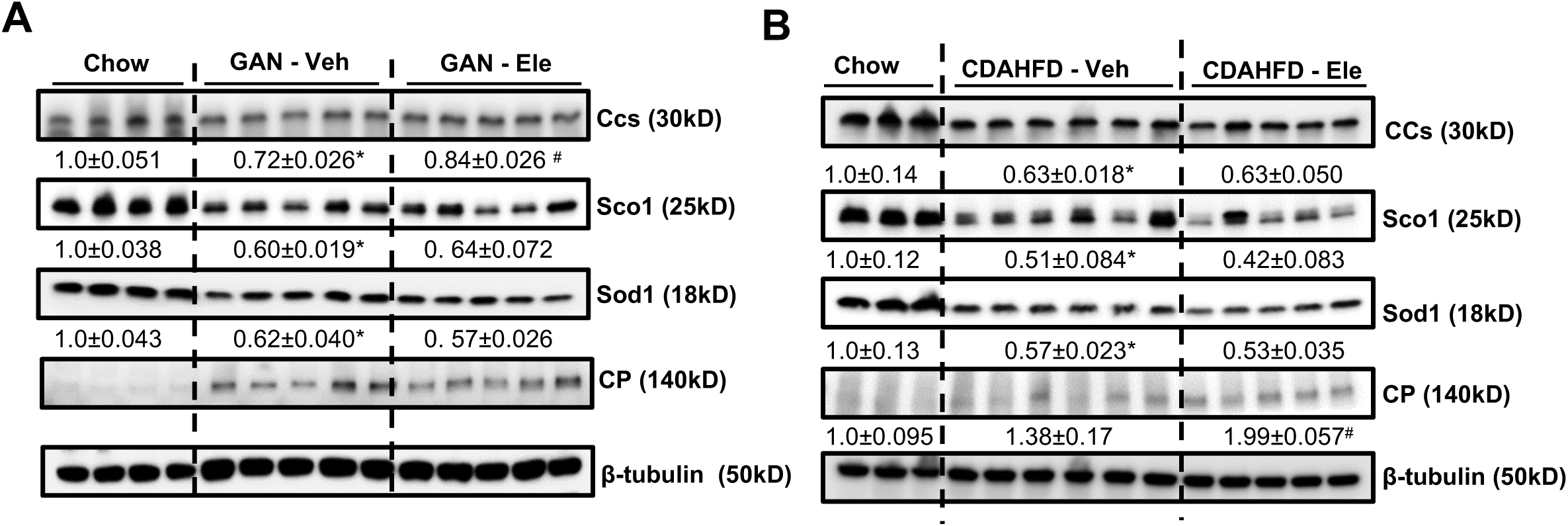
Restoring mitochondrial copper homeostasis improves mitochondrial function in MASH. (A, **B)** Immunoblot analysis of copper-dependent proteins in livers from chow-, GAN-, and CDAHFD-fed mice treated with vehicle or elesclomol. β-tubulin was used as a loading control. Densitometric quantification is shown below each blot. Data are shown as mean ± SEM; \**p* < 0.05 vs. chow; ^#^*p* < 0.05 vs. vehicle.

**Supplementary Fig. 5.**
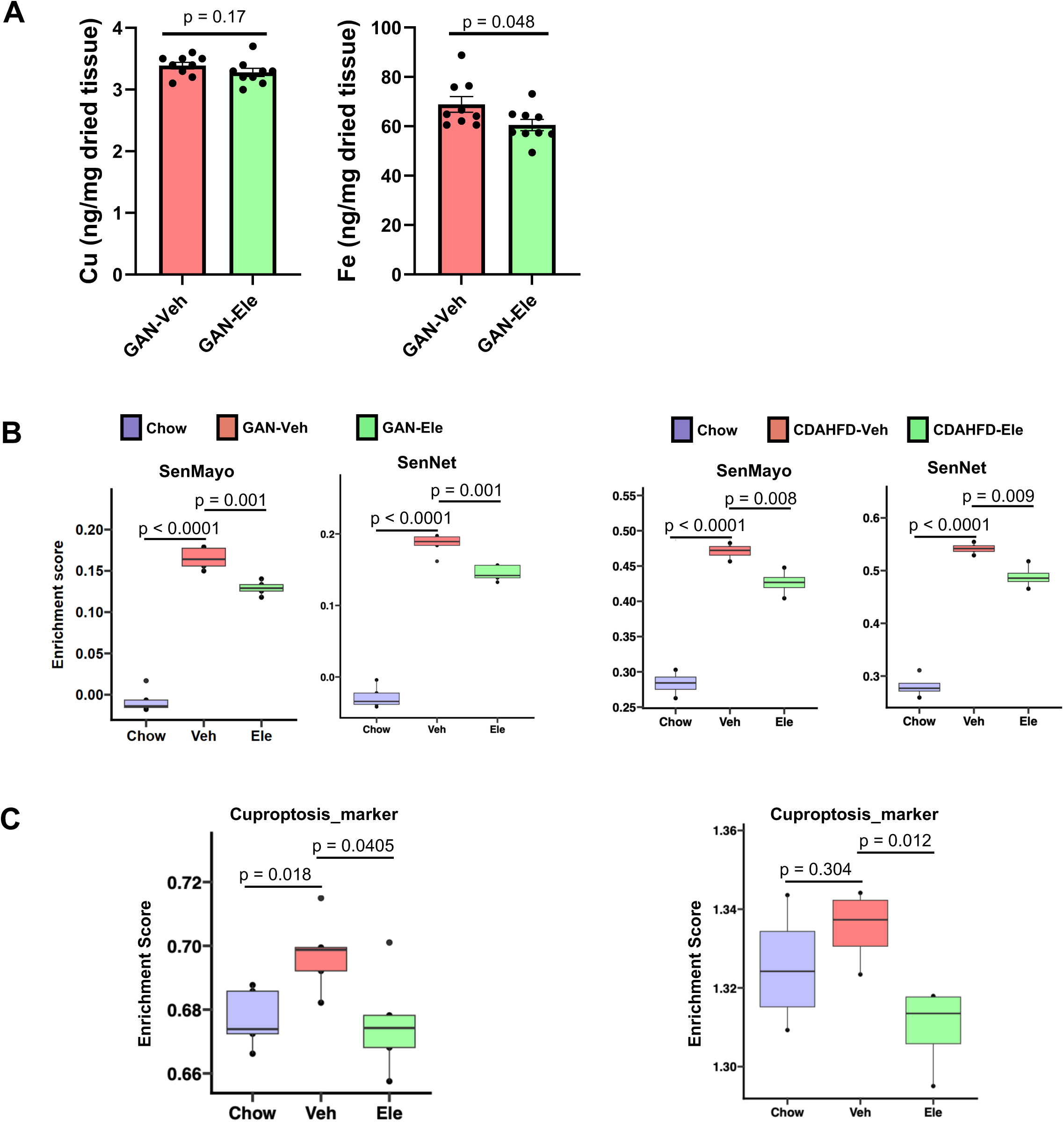
Restoring mitochondrial copper homeostasis suppresses hepatic iron accumulation and senescence without inducing cuproptosis. **(A)** Hepatic copper (Cu) and iron (Fe) levels in GAN-fed mice treated with vehicle or elesclomol, normalized to dried tissue weight. Data are shown as mean ± SEM**. (B)** Enrichment scores for senescence-associated gene signatures (SenMayo and SenNet) in livers from chow-, GAN-, and CDAHFD-fed mice treated with vehicle or elesclomol. **(C)** Enrichment scores for cuproptosis marker gene in chow-, GAN-, and CDAHFD-fed mice treated with vehicle or elesclomol. Enrichment scores were calculated using GSVA; data are shown as box-and-whisker plots with individual data points; statistical comparisons were performed using Wilcoxon or *t* tests as appropriate, with *P* values shown.

**Supplementary Fig. 6.**
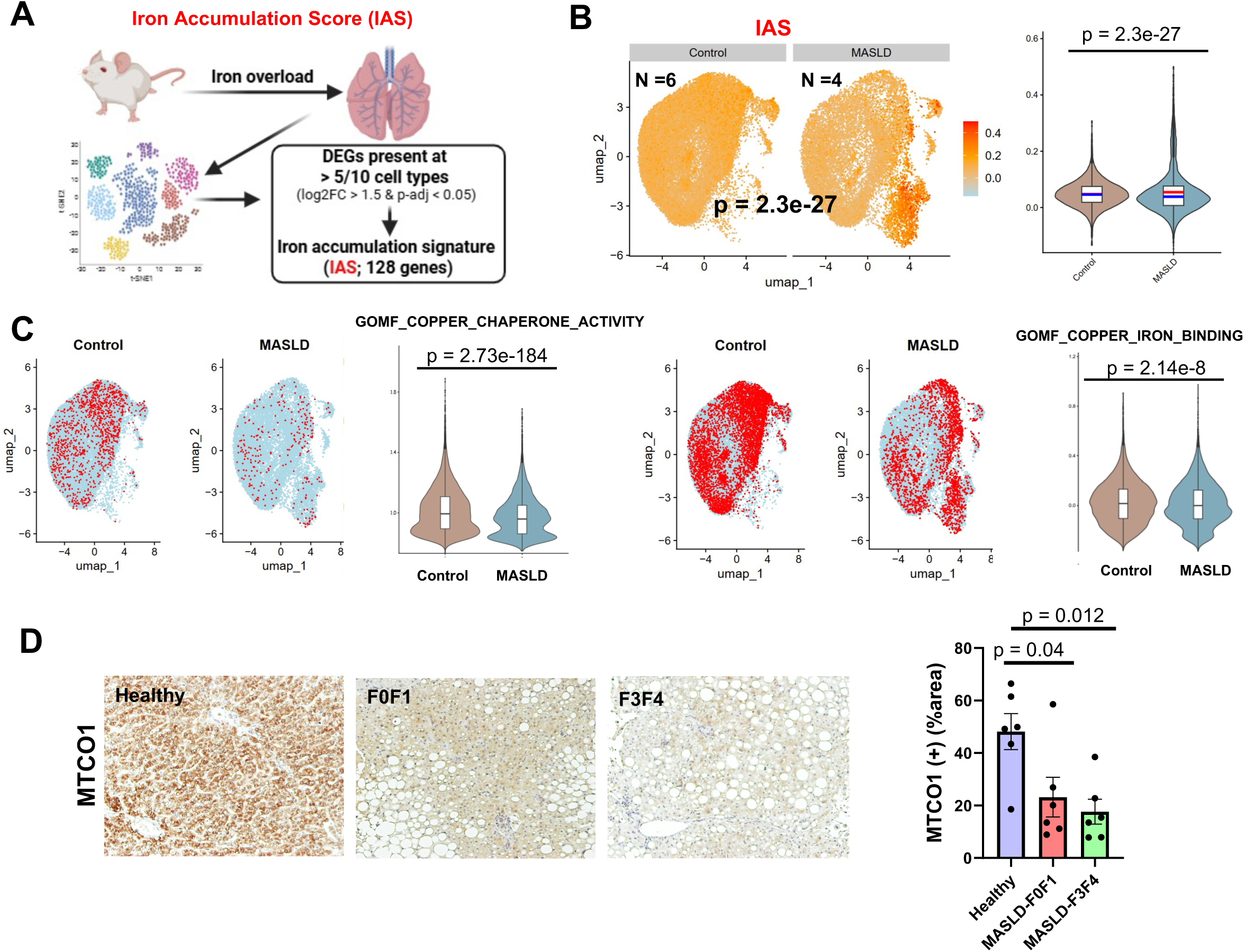
Copper homeostasis is dysregulated in hepatocytes of MASLD patients. **(A)** Schematic illustrating derivation of the Iron Accumulation Score (IAS) from iron-associated differentially expressed genes across multiple cell types. **(B)** Single-cell RNA-seq visualization of IAS enrichment in hepatocytes from control and MASLD livers, with corresponding violin plots showing IAS distribution. **(C)** Single-cell RNA-seq feature plots and violin plots showing enrichment of copper-related pathways, including copper chaperone activity and copper ion binding, in hepatocytes from control and MASLD livers. **(D)** Representative immunohistochemical staining for MT-CO1 in healthy control, F0-F1, and F3-F4 MASLD liver biopsies, with quantification of MTCO1-positive area. Data are shown as mean ± SEM; statistical comparisons were performed using one-way ANOVA with post hoc testing as indicated; *P* values are shown.

**Supplementary Fig. 7.**
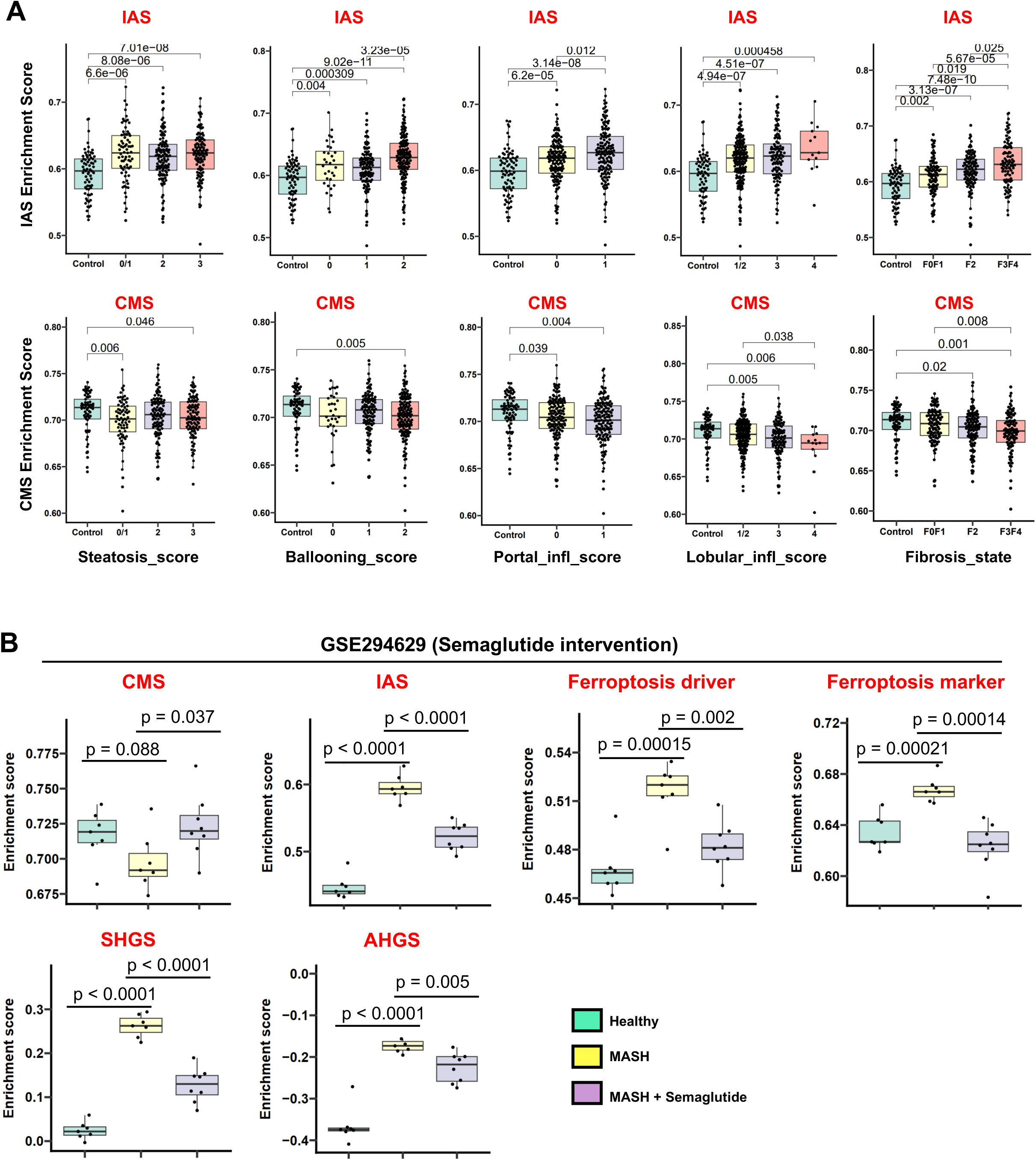
Copper–iron imbalance tracks MASLD progression, regression, and clinical outcomes. **(A)** CMS and IAS enrichment scores across histologic features of MASLD severity, including steatosis grade, hepatocellular ballooning, portal inflammation, lobular inflammation, and fibrosis stage, shown as box-and-whisker plots with individual data points. **(B)** CMS, IAS, ferroptosis driver, ferroptosis marker, SHGS, and AHGS enrichment scores in liver samples from healthy controls, MASLD patients, and MASLD patients treated with semaglutide (GSE294629). Enrichment scores were calculated using GSVA. Statistical comparisons were performed using one-way ANOVA or Wilcoxon tests as indicated; *P* values are shown.

**Supplementary Fig. 8.**
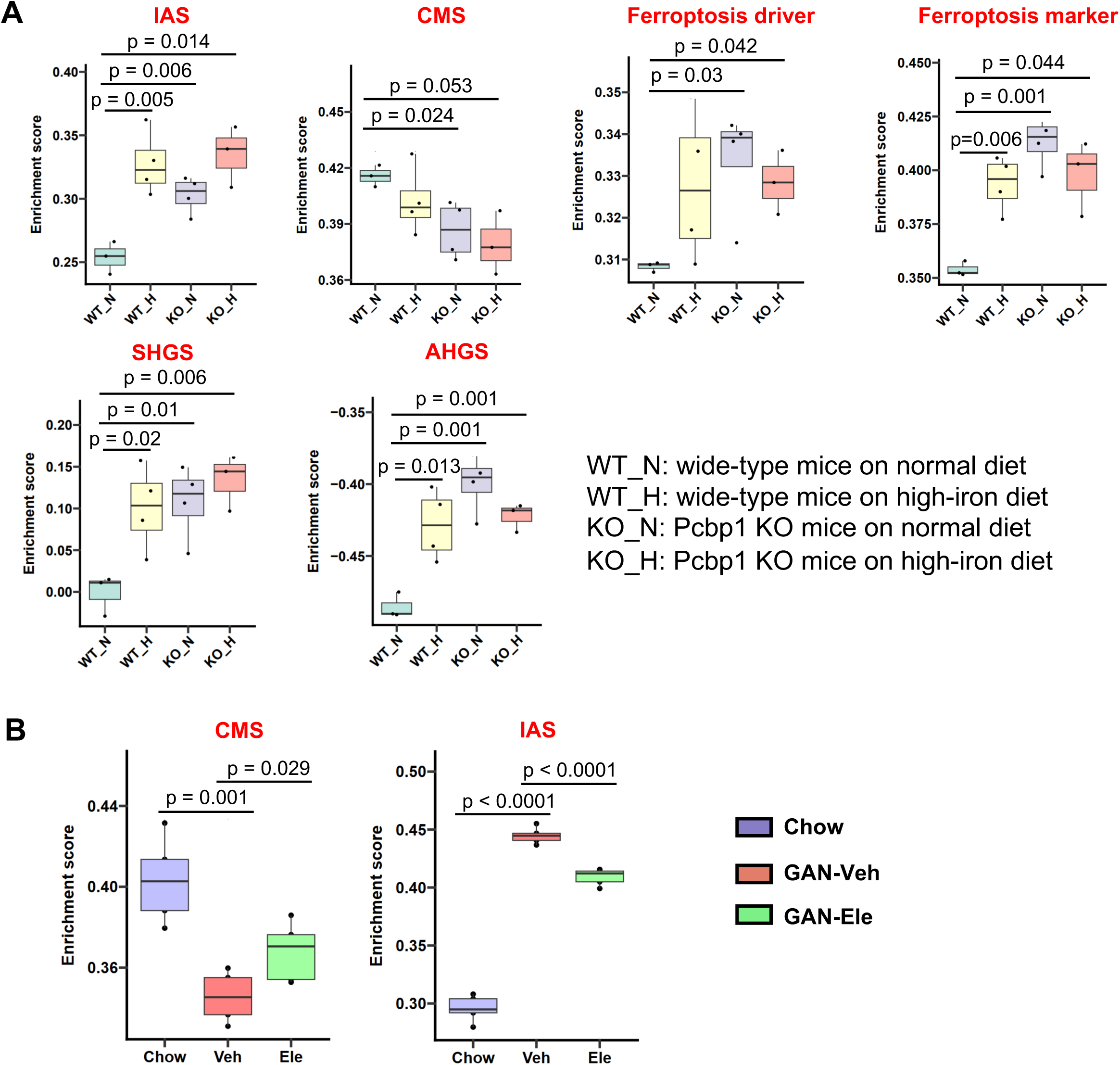
Copper–iron imbalance induces ferroptosis and senescence and is restored by elesclomol treatment. **(A)** Enrichment scores for Iron Accumulation Score (IAS), Cu-Mito Score (CMS), ferroptosis driver, ferroptosis marker, SHGS, and AHGS in livers from wild-type (WT) and Pcbp1 knockout (KO) mice fed a normal diet (N) or high-iron diet (H) from project PRJNA562934. Data were shown as box-and-whisker plots with individual data points. **(B)** CMS and IAS enrichment scores in livers from chow-fed mice and GAN diet-fed mice treated with vehicle or elesclomol. Enrichment scores were calculated using GSVA. Statistical comparisons were performed using one-way ANOVA or Wilcoxon tests as indicated; *P* values are shown.

**Supplementary Fig. 9.**
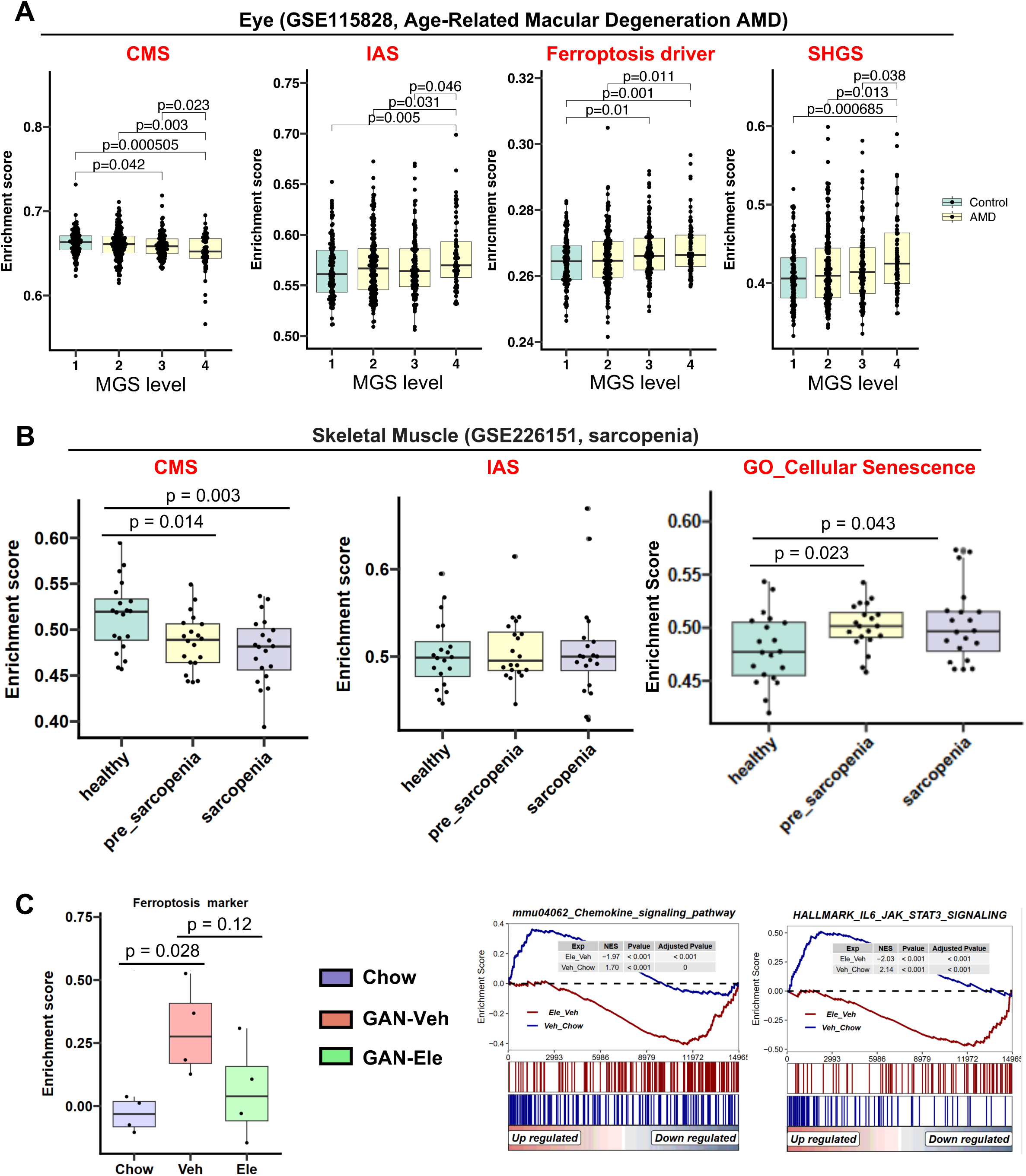
Copper-iron imbalance tracks MASLD-associated comorbidities, and restoring copper homeostasis suppresses kidney inflammation and fibrosis in MASH mice. **(A)** CMS, IAS, ferroptosis driver signature, and SHGS enrichment scores in retinal tissue from control subjects (MGS level of 1) and patients with age-related macular degeneration (AMD, MGS >1; GSE115828). **(B)** CMS, IAS, and cellular senescence gene ontology (GO) enrichment scores in skeletal muscle from healthy controls, pre-sarcopenia, and sarcopenia patients (GSE226151). **(C)** Enrichment scores for ferroptosis marker genes and GSEA plots for chemokine signaling and IL-6–JAK–STAT3 signaling pathways in kidneys from chow-fed mice and GAN diet-fed MASH mice treated with vehicle or elesclomol. Enrichment scores were calculated using GSVA; data are shown as box-and-whisker plots with individual data points. Statistical comparisons were performed using one-way ANOVA or Wilcoxon tests as indicated; *P* values are shown.

**Supplementary Table 1.**
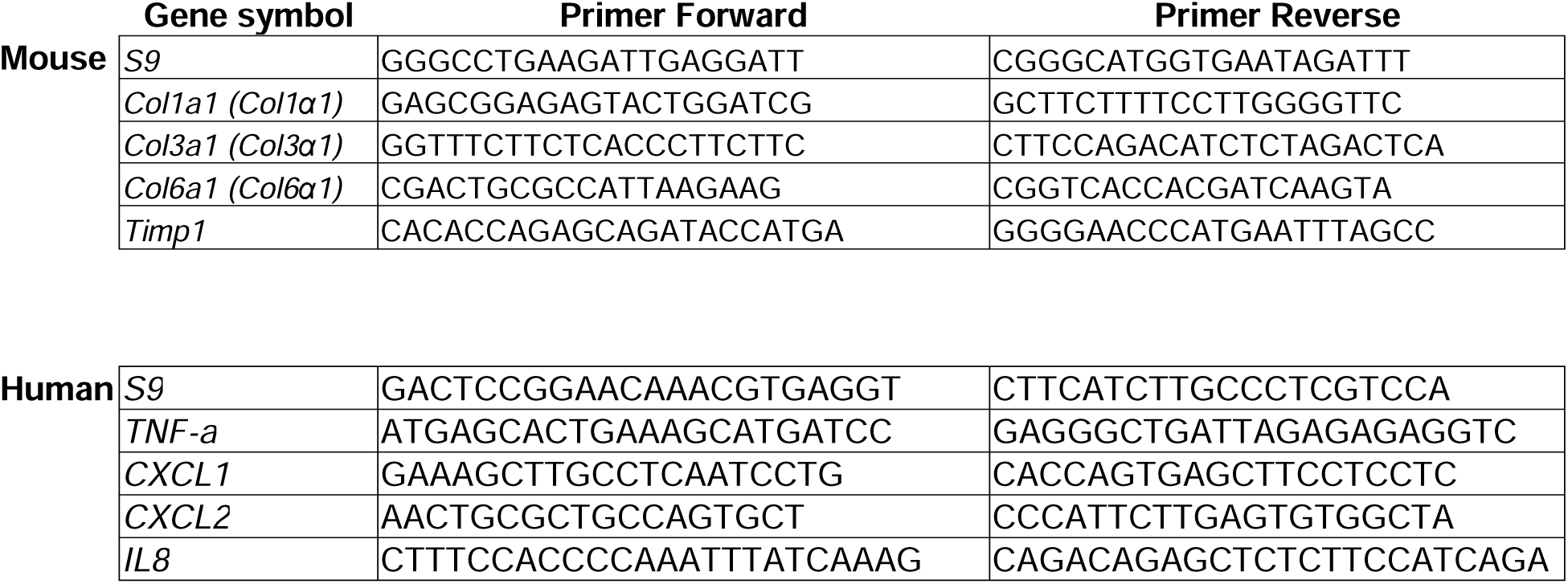
Primer used for qRT-PCR.

**Supplemental Table 2:**
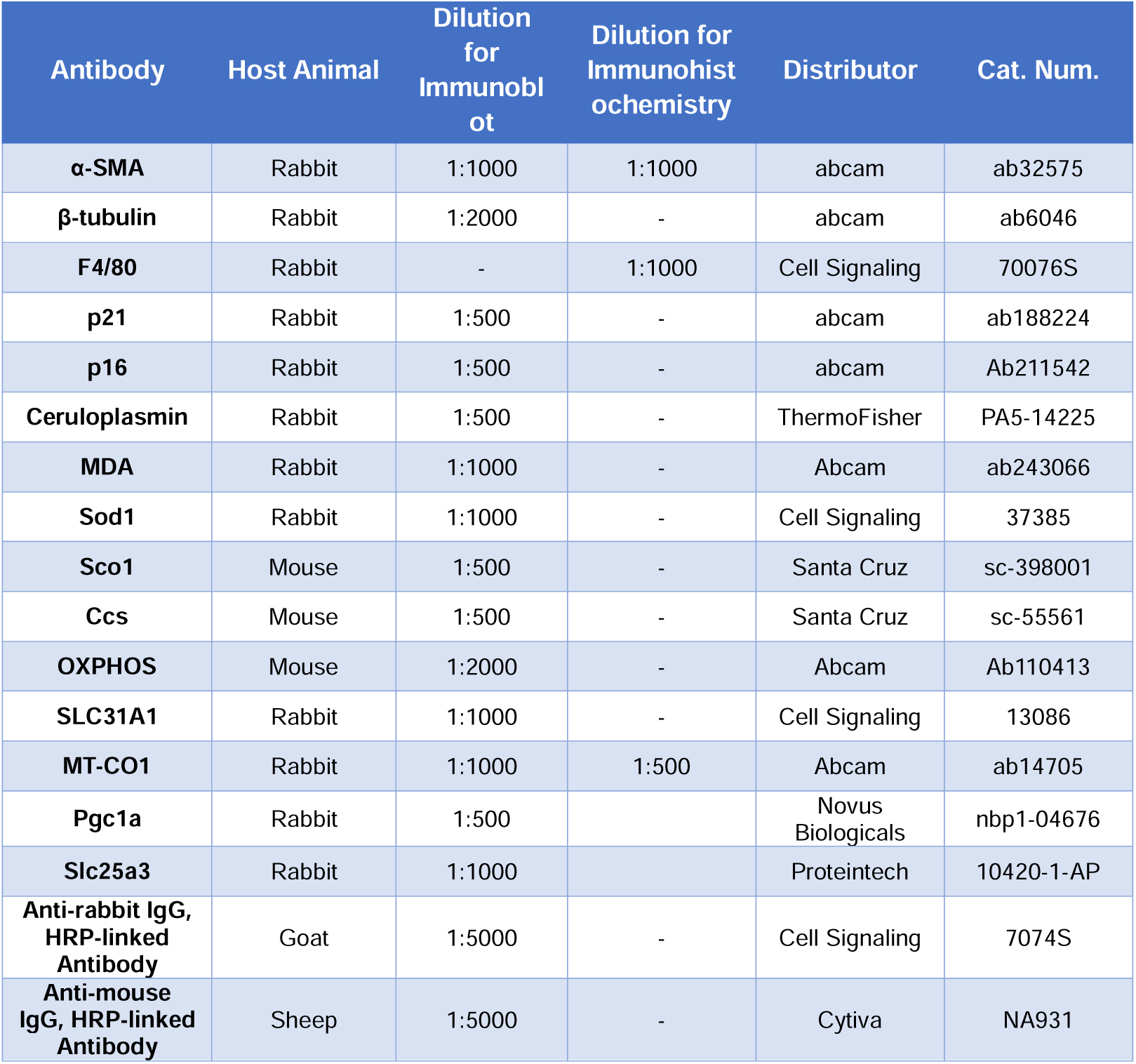
Primary antibody used for immuno-blot, histochemistry.

